# Small Extracellular Vesicles from Radioresistant H3K27M-Pediatric Diffuse Midline Glioma Cells Modulate Tumor Phenotypes and Radiation Response

**DOI:** 10.1101/2025.04.14.648723

**Authors:** Viral D. Oza, Kenan A. Flores, Yelena Chernyavskaya, Majd A. Al-Hamaly, Caitlyn B. Smith, Ronald C. Bruntz, Jessica S. Blackburn

## Abstract

Pediatric diffuse midline gliomas with the Histone 3 lysine 27-to-methionine mutation (H3K27M-pDMG) are aggressive brain tumors characterized by intrinsic resistance to radiation therapy, the current standard of care. These tumors exhibit significant intratumoral heterogeneity, with distinct subclonal populations likely contributing to therapy resistance. Emerging evidence suggests that small extracellular vesicles (sEV) mediate oncogenic signaling within glioma stem cell populations, yet their role under radiation-induced stress remains poorly understood. In this study, we characterized sEV uptake dynamics among H3K27M-pDMG tumor cells, identified key sEV surface proteins, and demonstrated that sEVs derived from radioresistant (RR) H3K27M-pDMG cells confer radioprotective effects on radiosensitive tumor cells. Molecular profiling revealed that RR-sEVs carry proteins, microRNAs (miRNAs), and metabolites associated with glycolysis, oxidative phosphorylation, and DNA repair. Upon uptake, RR-sEVs reprogrammed recipient cells by altering gene expression and metabolic pathways, and enhancing DNA repair and survival following radiation exposure. These findings provide insights into the role of sEV-mediated intratumoral communication as a contributor to radiation resistance in H3K27M-pDMG and suggest potential therapeutic strategies to disrupt this process and enhance radiation efficacy.

## 1 Introduction

Pediatric diffuse midline gliomas harboring the H3K27M mutation (H3K27M-pDMG) are aggressive, high-grade brain tumors and the leading cause of brain tumor-related deaths in children^1^. These tumors are characterized by resistance to radiotherapy, the current standard of care, resulting in a dismal five-year survival rate of less than 2%^2^. Surgical resection is nearly impossible^3^, and the tumor’s significant intratumoral heterogeneity drives diverse mechanisms of therapy resistance^4^. Despite over 95 clinical trials focused on H3K27M-pDMG, no effective therapies have emerged^5^, indicating an urgent need for a deeper understanding of the tumor’s biology and resistance mechanisms.

Radiation resistance in H3K27M-pDMG has been attributed to the dysregulation of pathways involved in DNA damage repair, cell cycle progression, chromatin remodeling, and key signaling cascades such as mitogen-activated protein kinase (MAPK), phosphoinositide 3-kinase (PI3K) and receptor tyrosine kinase (RTK) signaling pathways^6^. Mutations in *TP53* further contribute to radioresistance phenotypes^7,8^.However, much of what is known about radiation resistance in these tumors has focused on cell-intrinsic properties, while relatively little is understood about how interactions within the tumor microenvironment may also drive therapy resistance. Given the extensive intratumoral heterogeneity of H3K27M-pDMG^9^, it is possible that intrinsically radioresistant subclones could support the survival of radiosensitive cells within the same tumor, allowing the tumor to persist despite treatment.

Subclonal populations can communicate with and influence neighboring cells via paracrine signaling^10,11^ and small extracellular vesicles (sEVs)^12^. These vesicles, which carry proteins and nucleic acids, are known to promote oncogenesis in adult gliomas through the transfer of various oncogenic cargo^13,14^. In pediatric glioma, sEVs influence neural stem cell gene expression by downregulating genes involved in tumor suppression, differentiation, and the cell cycle^15^. Importantly, sEV cargo, such as microRNAs (miRNAs), can have subclonal specificity^12^, with distinct molecular signatures that can alter tumor behavior, such as invasive and migratory properties^15,16^. Despite these findings, the role of sEVs in radiation resistance in H3K27M-pDMG remains largely unexplored.

Radiation-induced stress alters sEV cargo in other cancers, such as head and neck and prostate cancer^17^, where sEVs carrying oncogenic proteins or interleukin-8 enhance radioresistance in recipient cells^18^. In adult gliomas, sEVs collected after radiation-induced stress have been shown to increase the survival of glioma cells by carrying altered cargo^19^. In healthy brain tissue, sEVs derived from oligodendrocytes, thought to be the precursors of H3K27M-pDMG, support neuronal metabolism and axonal transport^20^, suggesting a broader functional role for sEVs in cell survival and adaptation. Despite this, no studies have directly examined whether sEVs from radioresistant cells can enhance the survival of radiosensitive tumor cells in pediatric gliomas.

In this study, we investigate whether sEVs from radioresistant (RR) H3K27M-pDMG cells promote radiation resistance in radiosensitive (RS) pDMG cells. Using a treatment-naïve model, we demonstrated that RR-sEVs confer radioprotective effects through multiple mechanisms. We characterized H3K27M-pDMG sEV uptake dynamics and found that cell uptake is receptor-dependent. Molecular profiling revealed oncogenic miRNA, glycolytic and integrin-associated proteins, and numerous small metabolic molecules enriched in RR-sEVs. Functional experiments using biosensors showed that RR-sEVs induced rapid metabolic reprogramming, altered gene expression, and enhanced DNA repair in RS cells, ultimately increasing their survival and delaying radiation-induced cell death. Collectively, these findings provide the first detailed evidence of sEV-mediated cell-cell communication driving radioresistance in H3K27M-pDMG and identify novel therapeutic targets to disrupt these interactions.

## 2 Methods

### 2.1 Cell Culture and Maintenance

H3K27M-pDMG cell lines (SF7761 and SF8628) were obtained from Millipore Sigma. SF7761 cells were cultured as three-dimensional neurospheres in ReNcell Maintenance Media (SCM005, EMD Millipore) supplemented with bFGF (20 ng/mL; SRP3043, Sigma), EGF (20 ng/mL; GF144, EMD Millipore) and Penicillin-Streptomycin-Glutamine (1X; 10378016, Gibco). SF8628 cells were cultured in DMEM-High Glucose (D6546, Sigma) with 10% FBS (S11150H, R&D Systems), and 1X Penicillin-Streptomycin-Glutamine. PBT lines (gift from Nicholas Vitanza) were grown in Neurocult NS-A human proliferation kit (05751, VWR) with glutamax (1X; 35050061 Fisher Scientific), EGF (40 ng/mL), and bFGF (40 ng/mL). Plates were coated overnight with laminin (1:100 dilution in PBS; L2020, Sigma) at 4°C. SU-DIPG XIII cells (gift from Michele Monje) were cultured in tumor stem media consisting of 1:1 Neurobasal A (10999022, Fisher Scientific) and DMEM/F12 (11320033, Life Technologies), B27 (17504044, Life Technologies), bFGF, EGF, PDGF-A/B (10787-398 and10787, VWR), and heparin (10 ng/mL; S1346, Selleck Chemicals). BCH869 cells were cultured as per vendor instructions (SCC218, Sigma).

For passaging, adherent cells were lifted with Accutase (A7089, Sigma) or Trypsin-EDTA (25300062, Life Technologies). Neurospheres were dissociated with TrypLE Express Enzyme (12604021, Life Technologies) and DNase I (1:1000; 10104159001, Sigma). Briefly, for all suspension cell lines, cells were pelleted and resuspended in 1:1000 DNase in 2mL of TrypLE, incubated a nutator at 37°C for 10 minutes, and quenched with HSBSS (55037C, Sigma). Cells were pelleted at 300 × g for 7 minutes, then resuspended in cell culture media. All cells were maintained at 37°C with 5% CO_2_, routinely tested for mycoplasma, and used within 10 passages for experiments.

### 2.2 Clonal Isolation via Limiting Serial Dilution

Serial dilution was performed in ultra-low attachment microplates (7007, Corning). 200 µL of a cell suspension of 2x10^4^ cells/mL was deposited into well A1, then serially diluted 1:2 across the remaining wells, with a 100 µL media added to each well. Clones were detected using brightfield microscopy after 4 to 5 days. Clones were expanded from wells with a plating probability of <1 cell/well and subcultured into progressively larger plates over 1-4 weeks.

### 2.3 Radiation Treatment

Cells in 96-well plates were irradiated using a CS-137 gamma irradiator at 4.82 Gy/min on a rotating plate holder to ensure uniform dose distribution.

### 2.4 Generation of Reporter Stable Lines

To generate an SF7761 GEDI reporter line, the pMe:GC150-p2A-mApple construct (gift from the Finkbeiner Lab, Gladstone Institutes) was cloned into a pLenti-CMV-Puro destination vector (17452, Addgene) using Invitrogen Gateway technology (11791020, Life Technologies). Lentivirus was produced by transfecting HEK293T cells (12022001, Sigma) with the plasmid, psPAX2 and pMD2.g enveloping plasmids (12260 and 12259, Addgene) using TransIT reagent (2300, Mirus Bio) at a 1:0.1:0.9 ratio. Viral supernatant was collected at 48 and 72 hours post-transfection, concentrated using Lenti-X (631231, Takara Bio), and stored at -80°C. SF7761 cells were transduced with GEDI lentivirus and sorted for mApple-positive, GC150-low cells using an iCyt-Sony Cell Sorter. These cells were expanded and used for functional assays. For the SF7761 53BP1 reporter line, the mApple-53BP1trunc plasmid (69531, Addgene) was packaged into lentivirus as described above. Transduced cells were selected with puromycin (ant-pr-1, VWR) for 5-7 days and expanded for experiments.

### 2.5 Preparation of Concentrated Conditioned Media

Concentrated conditioned media (CCM) was prepared by culturing 6 x10^6^ cells in either 100mm dishes or T75 flasks. For SF8628 cells, conditioned media was harvested every 24 hours for 72 hours in media containing EV-depleted FBS (prepared by ultracentrifugation of FBS at 100,000 × g for 18 hours). Media was pooled and concentrated using Amicon-Ultra-15 filtration units with a 3 kDa cutoff (UFC900324, Millipore) by centrifuging 15 mL of media to a final volume of 1.5 mL at 2,000 × g at 4°C for 30 minutes. For SF7761 cells, conditioned media was collected every 3-4 days for 2 two weeks, pooled, and concentrated similarly.

### 2.6 Small Extracellular Vesicle (sEV) Isolation

sEVs were isolated via differential centrifugation according to MISEV 2023 guidelines^21^. Cells were grown at 37°C in 5% CO_2_ in appropriate culture conditions: neurospheres in T75 flasks for SU-DIPG XIII, SF7761, SCC218, and SF7761; two-dimensional adherent cultures in 150 mm dishes for SF8628; laminin-coated T75 flasks for PBT-22 and PBT-27 cells. Cell viability was verified to exceed 95% via trypan blue staining. Conditioned media (CM) was harvested based on cell type: every 24 hours for adherent cultures and every 96 hours for neurosphere cultures.

CM was centrifuged sequentially at 300 × g for 5 minutes (remove cells), 2,000 × g for 20 minutes (remove debris), and 10,000 × g for 40 minutes (remove large particles). Supernatants were filtered through a 0.22 µm filter (SLGL0250S, Sigma) to remove >220 nm particles, then spun at 100,000 × g for 90 minutes at 4°C. Pellets were washed in 0.22 µm filtered 1X PBS, centrifuged at 100,000 × g for 70 minutes at 4°C, and resuspended in 150 µL filtered PBS. Final sEV suspensions were aliquoted and stored at -80°C until use. sEVs were spin-filtered through a 0.5 mL 100 kDa Amicon filter (UFC510008, Millipore) prior to experiments. Flow-through solutions served as controls.

### 2.7 Nanoparticle Tracking Analysis

Isolated sEVs were resuspended in PBS and analyzed for size and concentration using a Zetaview (ParticleMetrix). Measurements were optimized to a particle concentration of 100-200 particles/frame. Camera settings included sensitivity:150-250, shutter:74-76, and cell temperature: 25°C. Acquired videos were analyzed by Zetaview software version 8.05.12. sEV concentrations used for functional studies were normalized to the number of particles per cell.

### 2.8 Transmission Electron Microscopy (TEM)

Samples were prepared for negative staining as described in Jung et al^22^. Briefly, formvar/carbon-film-coated 300-mesh copper EM grids (100502-760, VWR) were glow discharged for 1 minute. sEVs were fixed in 2% paraformaldehyde for 5 minutes, after which 5μL of sEV suspension was loaded onto a grid and incubated for 1 minute. The grid was exposed to 20 drops of 1% uranyl acetate solution using a syringe. Following staining, the grid was rinsed with a single drop of water and dried for 10 minutes inside a chemical hood. TEM was performed at the Electron Microscopy Center at the University of Kentucky using a FEI Talos F200X operating at 80keV, with a spot size of 6 and a 2-second exposure time.

### 2.9 sEV Protein Quantification and Immunoblotting

sEVs were lysed with RIPA buffer (P189900, VWR) and Halt Protease Inhibitor (1:100; 76466-056, VWR) on ice. Lysates were sonicated for 30 seconds and incubated on an orbital shaker at 4°C for 15 minutes. Protein content was quantified using a Pierce Micro BCA Protein Assay Kit (PI23235, VWR) with 1:100 diluted samples according to the manufacturer’s directions.

For immunoblotting, samples were mixed with 4X Laemelli buffer (1610747, Bio-Rad) containing ß-mercapto-ethanol (76177-226, VWR) to a final concentration of 1X, heated to 85°C for 5 minutes and cooled on ice for 3 minutes. Proteins were separated on 4-20% Min-PROTEAN TGX Stain-Free Gels (4568094, Bio-Rad) at 150V for 55 minutes and transferred to an Immuno-Blot PVDF Membrane (162055, Bio-rad) using the Trans-Blot Turbo Transfer System (Bio-rad) at 25V for 7 minutes. Membranes were blocked in 5% milk in 0.1% TBST for 1 hour and incubated overnight at 4°C with primary antibodies (1:1000 dilution) for Calnexin (ab22595, Abcam), HSP70 (EXOAB-Hsp70A-1, SBI), Histone H3 (ab1791, Abcam), CD9 (EXOAB-CD9A-1, SBI), CD63 (EXOAB-CD63A-1, SBI), and CD81 (EXOAB-CD81A-1, SBI) in a 5% milk with 0.1% TBST. Blots were washed three times with 0.1% TBST for 5 minutes and incubated for 1 hour at room temperature with goat anti-rabbit-HRP secondary antibody (1:1000 dilution; 20403, SBI). Bands were visualized using the Clarity Western ECL Substrate Kit (1705060, Bio-rad) on a ChemiDoc Touch Imaging System (Bio-Rad).

### 2.10 Colony-Forming Assay

Cells (2500-5000/well) were seeded in flat-bottom 96 well plates and imaged every 3-7 days using a Biotek Lionheart FX System. Media was replenished every 3-4 days. Colony images were processed with Gen5 software using rolling ball background subtraction and point spread deconvolution. Images were analyzed using a custom ImageJ script with manual thresholding for colony quantification.

### 2.11 Mitochondrial Stress Test via Seahorse Assay

A mitochondrial stress test was conducted using the Agilent Seahorse xe96 platform. Cells were plated at 50,000 cells/well on Cell-Tak (47743, VWR) coated seahorse plates (102978, Agilent). Two hours after cell seeding, the sensor cartridge was hydrated. Three successive injections of Oligomycin, FCCP (2 µM), and Rotenone/Antimycin were administered over the course of 90 minutes in DMEM assay medium without FBS. Data were analyzed using WAVE software, with normalization based on cell counts.

### 2.12 Caspase 3/7 Assay

sEVs at low (∼10^7^), medium (∼10^8^), and high (∼10^9^) concentrations were spiked into 100 µL of SF7761 media for 30 minutes. Caspase-Glo 3/7 Reagent (PAG8091, VWR) was added (100 µL/well) and incubated for 30 minutes. Luminescence was measured using a Synergy LX Multi-Mode Reader (Biotek). Media-only wells were used as blanks for normalization.

### 2.13 Cell Titer Glo Assay

For cell line and CM viability assay, cells were plated at a density of 5,000 cells/well in 96-well optical plates (29444, VWR) with 100 µL of media. After 4 hours, cells were irradiated and immediately transferred back to an incubator at 37°C with 5% CO_2_. For wells treated with CM, conditioned media was concentrated by 10X using a Amicon Ultra 15 (UFC900324, Millipore) before 100 µL of conditioned media was added for 24 hours before irradiation. After 72 hours post irradiation, plates were equilibrated to room temperature, and 100 µL of Cell Titer-Glo Luminescent Cell Viability Assay Reagent (PAG7572, VWR) diluted 1:3 in PBS was added to each well. Plates were shaken on an orbital shaker for 10 minutes. Luminescence was measured using a Synergy LX Multi-Mode Reader (Biotek) with media-only wells serving as blanks for normalization.

For ATP measurements in sEVs, (∼10^8^) sEVs were spiked into 100 µL of SF7761 media and incubated for 30 minutes. Cell Titer-Glo Reagent (100 µL) was added to each well, and luminescence was measured following the same protocol as above.

### 2.14 sEV Uptake and Flow Cytometry Analysis

sEVs were labeled with Vybrant CFDA SE Cell Tracer dye (V22885, Life Technologies). A total of 1-5 × 10^8^ sEVs were incubated with 40 µM CFDA at 37°C for 2 hours, concentrated and purified using an Amicon Ultra-0.5 Centrifugal Unit. Labeled sEVs were added to cells plated at 40,000 cells/well at a ratio of 2,500-5,000 sEVs/cell.

To assess the effects of inhibitors on sEV uptake, cells were pre-treated for 30 minutes with either Dynasore (80 µM; SML0340 Sigma), Heparin (100 µg/mL; S1346, Selleck), MBCD (5mM; C4555, Sigma), or Amiloride (100 µM; BML-CA200, Enzo Life Sciences). For proteinase protection assays, sEVs were treated with Proteinase K (100 µg/mL; CB3210-5, Denville Scientific) for 5 minutes at 37°C, followed by inactivation with 0.2 µL of boiled PMSF (329-98-6, Tocris) on ice for 10 minutes.

To prepare cells for flow cytometry, cells were washed in PBS, incubated with TrypLe Express for 10 minutes at 37°C, and quenched with 180 µL of HSBSS, and strained through a 40 µm cell strainer (21008-948, VWR) for flow cytometry analysis. Flow Cytometry was performed on a BD Symphony A3, with CFDA fluorescence measured in the BB15 channel. Positive signals were gated based on fluorescence intensity above the unstained cell gate, and additional gating was applied to distinguish CFDA-stained cells undergoing multiple cell divisions. Side scatter and forward scatter differentiated single cells from cell debris. A consistent number of events were captured for all samples, and data were analyzed using FlowJo v10.

### 2.15 sEV-Mediated Functional Changes in SF7761 Cells

SF7761 cells were plated in 96-well black wall plates at a density of 25,000 cells/mL (100 µL/well). After 4 hours, cells were treated with RR-sEVs, RS-Clone 1-sEVs, or control (100 kDa ultrafiltrate) at a concentration of 5,000 sEVs/cell for 18 hours. Reporter cells were imaged at 37°C with 5% CO_2_ and then irradiated. Imaging was performed at 6, 12, and 18 hours post-irradiation (hpi) for GEDI experiments, 0, 3, 9, 20 hpi for 53BP1 experiments, and 4 hpi for yH2AX immunofluorescence experiments.

### 2.16 Biosensor Assays for GEDI and 53BP1 Analysis

GEDI analysis was conducted using a Lionheart FX imager to capture 10 × 10 montages of wells in 96-well plates. Images were processed using Gen5 software for background correction using the point spread function followed by Gaussian deconvolution. Downstream analysis was performed using ImageJ. Masks were generated in the red channel to quantify fluorescence in the green and red channels using a custom ImageJ script. The GEDI threshold of cell death was determined by measuring the GEDI ratio of cells before and after a lethal dose of radiation (25Gy) at 24 hours. GEDI ratios were plotted over time using the tidyverse package in RStudio. Kaplan-Meier survival analysis classified cells as alive if they retained a GEDI ratio and as dead if they were no longer visible in the field of view (FOV).

For 53BP1 analysis, red-channel masks were generated using maximum contrast to detect foci with fluorescence intensities with at least an order of magnitude higher than background levels. The number of puncta per cell in the field of view was calculated using a custom script in Rstudio with the tidyverse package.

### 2.17 Immunofluorescence and Live/Fixed Cell Fluorescent Imaging

Cells were fixed in 96 well plates with 4% paraformaldehyde (102091-906, VWR), washed three times in 1X PBS, and permeabilized with 1%Triton X-100 (X100, Sigma) for 15 minutes at 4°C. Following three additional PBS washes, cells were blocked in 1% BSA (BP9706, Fisher Bioreagents). The primary antibody, Phospho-Histone H2A.X (9718, Cell Signaling Technologies) was applied at a 1:200 dilution overnight at 4°C. Cells were then washed with 1X PBST (PBS with 0.05% Tween 20; P1379, Sigma) and incubated for 1 hour with Alexa Fluor 488 goat anti-mouse IgG secondary antibody (1:1000; A11001, Invitrogen) and DAPI (1:1000; 62248, Life Technologies). After 3 final PBST washes, cells were imaged immediately or stored at 4°C for subsequent imaging.

Live and fixed cell imaging was performed using a Biotek Lionheart FX Automated Imaging System, capturing 10 × 10 montages per well at 20X magnification. GC150 and mApple reporters were imaged using Biotek GFP (PN:1225001,1225101) and RFP (PN:1225003,1225103) cube sets, respectively. Additional imaging was conducted using a Nikon A1R confocal imaging system. Image processing followed the workflow described for the colony-forming assay.

### 2.18 RNA Extraction, Library Preparation, and Next-Generation Sequencing

Total RNA was extracted from SF7761 cells and subclones using the Zymo Research Quick-RNA kit (R1054) and quantified using the Qubit RNA HS Assay Kit (Q10210, ThermoFisher). RNA integrity was assessed with an Agilent 2100 Bioanalyzer, and only samples with a RNA Integrity Number (RIN) greater than 9.5 were used for downstream applications. For mRNA enrichment, 250 ng of RNA was processed with the NEBNext Poly (A) mRNA Magnetic Isolation Module (E7490S, New England Biolabs) to selectively capture polyadenylated transcripts. Library preparation followed the NEBNext Ultra II Express Library Prep Kit for Illumina (E3330S, New England Biolabs) protocol, which included mRNA fragmentation at 94°C for 15 minutes, first- and second-strand cDNA synthesis, end-repair, adaptor ligation, and amplification using NEBNext Multiplex Oligos (E6448S, New England Biolabs). Library quality was assessed using an Agilent 2100 Bioanalyzer and concentrations were measured with the Qubit dsDNA HS Assay Kit. Equimolar library pools were sequenced on an Illumina NovaSeq 2500 in a 2 × 150bp configuration.

RNA was also extracted from purified and size selected sEVs using TRIzol Reagent (15596026, ThermoFisher) following the manufacturer’s instructions. Precipitation was performed overnight at -20°C using equal volume of isopropanol and Glycogen (0.5 µL; R0551, ThermoFisher). RNA pellets were washed twice with 70% ethanol, air-dried, and resuspended in 10 µL of nuclease free water. For RNA extracted from sEV-treated cells, media (100 µL) was transferred into tubes with 800 µL of TRIzol, and 100 µL of TRIzol was added directly to each well of the 96-well plate. Lysates were pooled with their corresponding media/TRIzol mixtures. These samples followed the same RNA extraction and quantification procedures as above and all samples were treated and processed in triplicate.

Libraries for RNA extracted from sEV treated SF7761 cells were prepared using the SMART-Seq mRNA LP Kit (with UMIs) (634762, Takara) according to manufacturer’s instructions. Briefly, 10 ng of RNA was used as input for each sample. After first-strand cDNA synthesis, libraries were ligated with unique molecular identifier (UMI)-containing adapters enriched with 17 cycles of PCR. Libraries were validated for size and concentration using a Bioanalyzer and gel electrophoresis, pooled at a final concentration of 60 nM in 24 µL, and sequenced using the NovaSeq platform.

For miRNA sequencing, 5 ng of sEV RNA and 1 ng of positive control included in the kit were used as input for library preparation with the SMARTer smRNA-Seq Kit (635029, Takara). according to the manufacturer’s instructions. Library purification was carried out using the NucleoSpin Gel and PCR Clean-Up kit (740611.50, Takara). Size of purified libraries were validated using gel electrophoresis and quantified with the Accugreen High Sensitivity dsDNA Quantification Kit (75845-718, Biotinum).

### 2.19 sEV Metabolomics

Metabolites from sEV samples were extracted by resuspending 10 µL of sEV suspension in 300 µL 80:20 acetonitrile (ACN): H_2_O and homogenizing with a Disruptor Genie (Scientific Industries) at 3000 rpm for 3 minutes. Proteins were precipitated at -20°C for 30 minutes, followed by centrifugation at 17,000 × g for 10 minutes at 4°C. Samples were snap-frozen in liquid nitrogen and stored at -80°C. All solvents were LC-MS grade from Fisher Scientific.

For hydrophobic interaction liquid chromatography (HILIC), thawed extracts were transferred to LC-MS autosampler vials. Polar metabolites were separated using a Waters Atlantis Premier BEH Z-HILIC column (2.5 mm, 2.1 × 100mm) on a Thermo Vanquish UPLC liquid chromatography (LC) system with a binary solvent system of 10 mM ammonium acetate in water, pH 9.8 (mobile phase A) and acetonitrile (mobile phase B). The flow rate was 0.33 µL/min and the LC gradient was 100% mobile phase B from 0-1.6 min, 70% by 6.2 min, 40% by 7.7 min, 30% by 8.3 min, and returned to 100% B by 10.3 min to re-equilibrate until 13.7 min. Reverse-phase LC-MS analysis was performed on dried ACN extracts reconstituted in 100 µL H_2_O with 0.1% formic acid, using an Agilent InfinityLab Poroshell 120 EC-18 column on a Thermo Vanquish UPLC. The gradient ramped from 3% mobile phase B, (0 min) to 40% B (10 min), 95% B (12 min), and back to 3% B for re-equilibration until 17 min.

Both HILIC and reverse-phase separations were analyzed on a Thermo Exploris 240 Orbitrap Mass Spectrometer. Spectra were acquired in positive and negative ionization modes with data-dependent MS/MS acquisition. Parameters included a 1 Da isolation window, stepped collision energies (20, 30, and 60) and 120k MS1 resolution. Data processing with Compound Discoverer (CD) (version 3.3.3.200, Thermo Scientific) included untargeted metabolomics workflows and filtering for Δmass (-5 to 5 ppm), reference ion [M+H]^+^ or [M-H]^-^ for positive and negative ionization, respectively, retention time thresholds (< 8.5 min for HILIC, <11.5 min for reverse-phase columns), peak rating > 6, and peak area >300% higher than blank samples in at least two sEV samples.

### 2.20 sEV Proteomics

sEV proteins were digested using the Preomics iST kit according to the manufacturer’s protocol. Protein inputs (5 µg) were lysed in 100 µL of buffer and incubated at 95°C for 10 minutes, followed by digestion at 37°C for 3 hours. Purified peptides were vacuum-dried, resuspended in 5% acetonitrile with 0.1% formic acid and analyzed on a Thermo Scientific™ Orbitrap Exploris 240 mass spectrometer paired with a Vanquish Neo UHPLC system.

Peptides were separated on an EASY-Spray HPLC column using a linear gradient from 2-55% mobile phase B over 90 minutes at flow rate of 0.3 µL/min. Data-dependent acquisition (DDA) parameters included a 60k MS1 resolution, scan range of 375-1200 m/z and 15k MS2 resolution with 2 m/z isolation and 26% normalized HCD collision energy. Proteomics data were analyzed using Thermo Scientific Proteome Discoverer (version 3.1) software with the SEQUEST search algorithm and the Homo Sapiens UniProt database (downloaded March 2024). Precursor mass tolerance was set to 10 ppm. Quantification used unique and razor peptides normalized to total peptide abundance.

### 2.21 Bioinformatics Pipelines for RNA, miRNA, and Proteomics

For bulk RNA sequencing, Fastq files were aligned to the human reference sequence using STAR aligner. Transcript read counts were generated using HTSeq, and normalization and differential gene expression analysis were conducted using DESeq2. Gene set enrichment analysis was done using GSEA software.

For small RNA sequencing, adaptors were trimmed using Cutadapt, and reads were mapped to mature and hairpin miRNA sequences using MiRDeep2. Quantification was performed and reads were normalized using the TMM method in EdgeR to generate CPM values. MiRNA-target interaction networks were constructed using Mienturnet, including only experimentally validated interactions.

Proteomic pathway analysis was conducted with PANTHER 19.0. Statistical overrepresentation analysis was performed using Fisher’s exact test with a false discovery rate (FDR) correction of 0.05. Functional classification was based on PANTHER Protein Class annotation data.

Metabolomics analysis focused on targeted TCA cycle pathways. One-factor statistical analysis was performed in MetaboAnalyst.

### 2.22 Statistical Analysis

Data are presented as mean +/- standard error (SEM). Statistical analyses were performed using Prism 8 (Graphpad software), GSEA software, MetaboAnalyst, and PANTHER. miRNA-target enrichment and network analysis were conducted in Mienturnet. Statistical tests included one-way ANOVA with Tukey’s multiple-comparison test and two-way ANOVA, depending on dataset and experimental design.

## 3 Results

### 3.1 Radioresistant subclones in a treatment-naïve H3K27M-pDMG cell line exhibit enhanced oxidative phosphorylation

H3K27M-pDMGs exhibit variable responses to radiation, likely driven by intratumoral heterogeneity and specific mutational profiles^23,24,25^. Intrinsic radioresistance, a characteristic feature of glioblastoma stem cells, may contribute to this variability^26^. To investigate whether clonal subpopulations within a treatment naïve (i.e. no prior radiation) radiosensitive (RS) H3K27M-pDMG cell line harbor intrinsic radioresistance, we isolated subclonal populations (RS-Clone 1, RS-Clone 4, RS-Clone 5, RS-Clone 6) from the parental SF7761 (RS) cell line (Figure 1A). We then compared these clones to SF8628, a K27M mutant pDMG cell line derived from a patient biopsy pre-treatment that shows strong radioresistance (RR)^27^.

**Figure 1:**
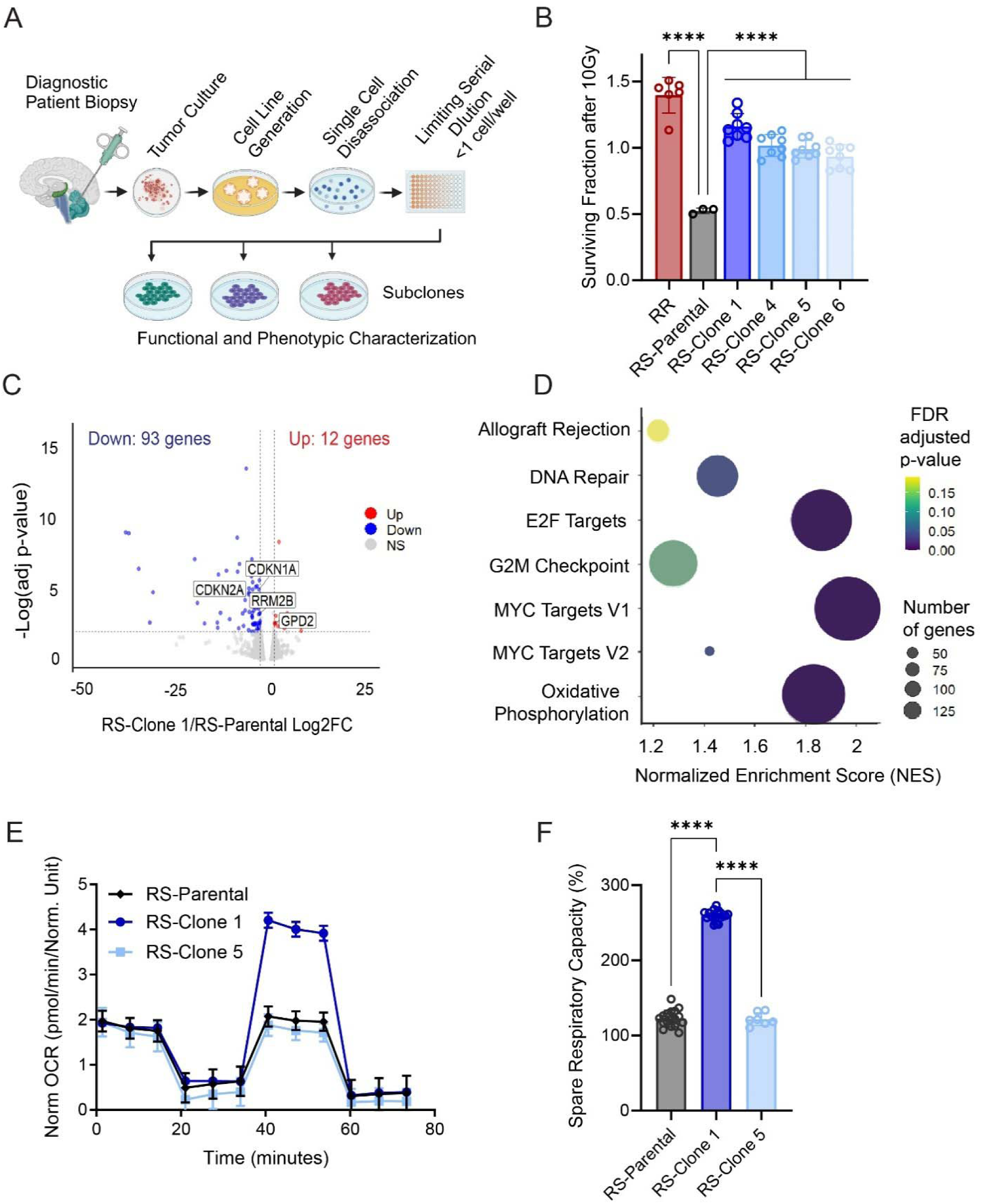
Clonal isolation from an H3K27M-pDMG bulk radiosensitive population reveals subclones with intrinsic radioresistant and distinct metabolic profiles. (**A**) Schematic representation of the clonal isolation process using limiting serial dilution from a bulk radiosensitive (RS) H3K27M-pDMG patient derived cell line (SF7761) (**B**) Cell viability using CellTiter-Glo in two treatment-naïve H3K27M-pDMG cell lines, SF7761 (RS), and SF8628 (a bulk radioresistant [RR]), along with four RS-derived subclones, at 72 hours post radiation dose (10 Gy) across two biological replicates. Values are normalized to 0 Gy-treated controls, n=2 biological replicates. (**C**) Volcano plot depicting differentially expressed genes in RS-Clone 1 compared to the parental RS-population, generated from RNA sequencing across three biological replicates. The horizontal dashed line represents an FDR threshold of 0.05, and the vertical dashed line indicate a log_2_ fold change of -2 and 2. (**D**) Gene set enrichment analysis (GSEA) of RS-Clone1 compared to RS, highlighting significantly enriched gene sets (FDR ≤ 0.25). Bubble size corresponds to the number of genes within each pathway, and color represents the FDR-adjusted p-value. (**E**) Mitochondrial stress test and (**F**) quantification of spare respiratory as percentage of base line in RS-Parental and two RS-derived sublcones using the Seahorse platform across two biological replicates. Data are presented as means ± standard error of the mean (SEM). Statistical significance was determined using two-way ANOVA for (B) and (E), DESeq2 for (C), and GSEA software for (D). \**p* < 0.05, \*\**p* < 0.01, \*\*\**p* < 0.001.

We assessed cell viability following 10 Gy irradiation. RS-Clone 1 showed significantly higher cell viability than the parental SF7761 line (*p*<0.001), resembling the radioresistant phenotype of SF8628 (Figure 1B). Under both standard and radiation stress conditions, RS-Clone 1 formed significantly larger colonies than other subclones (*p*<0.001; Figure S1A-B), indicating enhanced self-renewal capacity. To further assess long-term survival, subclones were irradiated up to 60 Gy over several weeks, reflecting extended clinical exposure. RS-Clone 1 was the only clone to retain standard neurosphere morphology (data not shown).

To explore the molecular basis of RS-Clone 1’s phenotype, we performed RNA sequencing of the parental SF7761 (RS) line and RS-Clone 1. We identified 105 differentially expressed genes in RS-Clone 1 compared to SF7761 (Figure 1C), including those implicated in glycolysis, cell cycle progression, and DNA repair. Gene set enrichment analysis (GSEA) revealed significant enrichment of pathways regulating E2F targets (NES:1.86; FDR: 0), MYC targets (NES:1.96; FDR:0), oxidative phosphorylation (NES:1.96; FDR:0), and DNA repair pathways (NES:1.44; FDR:.03) (Figure 1D). Given RS-Clone 1’s transcriptional signature, we next examined mitochondrial metabolism. Metabolic profiling using Seahorse assays confirmed that

RS-Clone 1 had significantly higher spare respiratory capacity and maximal respiration compared to the parent (RS) SF7761 line and RS-Clone 5 (*p*<0.001) (Figures 1E-F). The RR cell line SF8628 exhibited the highest metabolic activity across all parameters (Figure S1C).

Together these data demonstrate that subclonal populations within a treatment naïve, radiosensitive H3K27M-pDMG cell line can harbor intrinsic radioresistance. Elevated oxidative phosphorylation is a feature of the radioresistant phenotype, suggesting a link between mitochondrial metabolism and radioresistance in these cells.

### 3.2 Conditioned media from a radioresistant line confers radioresistance to a radiosensitive H3K27M-pDMG and reveals common sEV features

Previous studies suggest that clonal H3K27M-pDMG cell secretomes can influence stem-like phenotypes in tumor cells^10^. To determine whether secreted factors impact radiation-induced cell survival, we treated radiosensitive (RS) SF7761 cells with concentrated conditioned media (CCM) from RS-SF7761 cells, RS-Clone 1, or radioresistant SF8628 (RR) cells for 24 hours prior to irradiation (4 Gy or 8Gy). Cell viability was assessed 72 hours post-irradiation and normalized to unirradiated (0 Gy) controls. CCM derived from RR-SF8628 significantly enhanced the viability of RS cells compared to both RS-SF7761 CCM (*p* <0.001) and RS-Clone 1 CCM (*p* <0.01); (Figure 2A). RS-Clone 1-CCM did not significantly alter viability compared to the parental SF7761 CCM, though a mild trend was observed. These findings indicate that fully radioresistant pDMG cells can promote radiation survival in radiosensitive cells, while partially resistant subclones may not exhibit the same strong effect.

**Figure 2:**
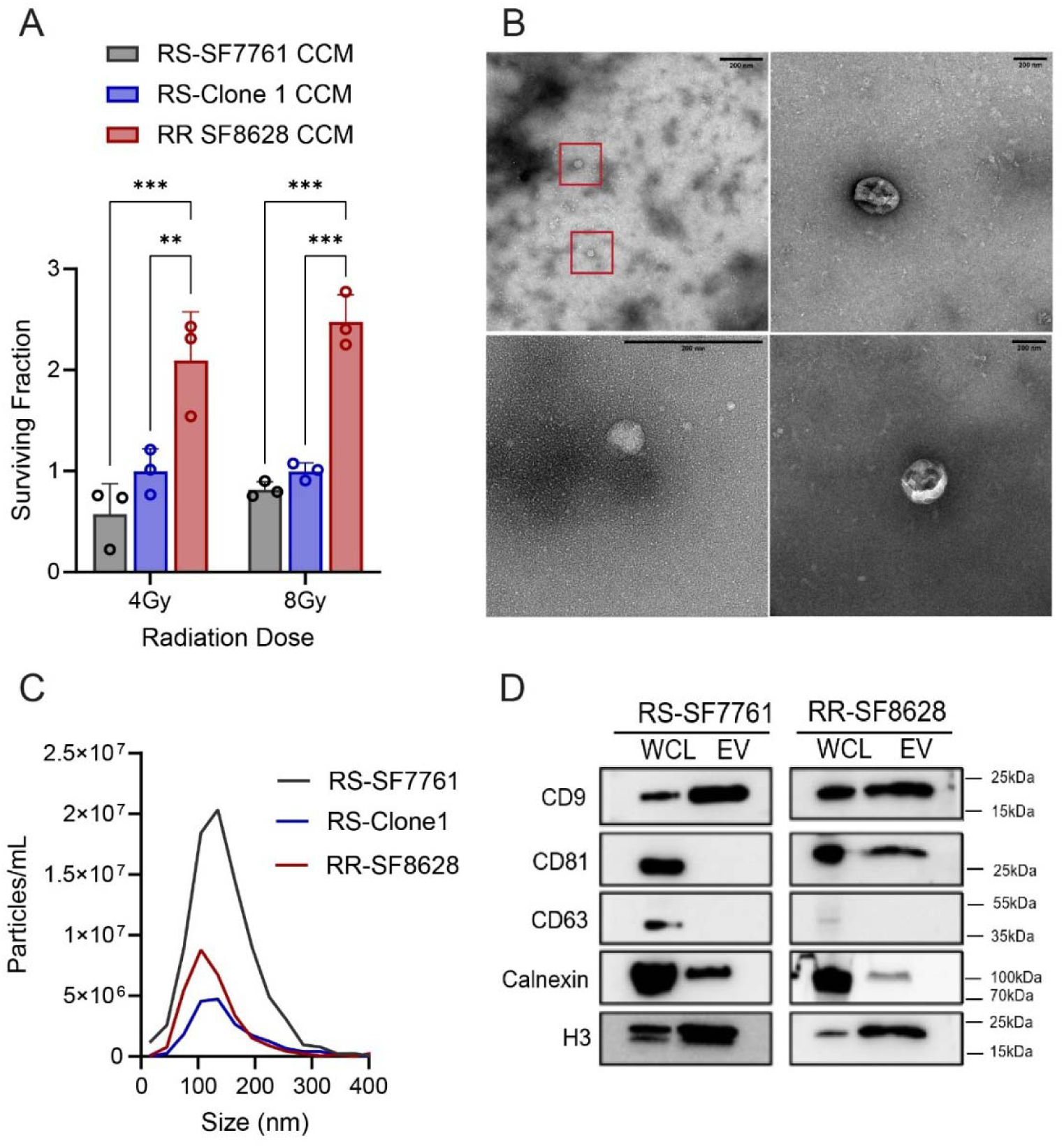
Conditioned media from radioresistant (RR) H3K27M-pDMG enhances radioresistance in RS cells and contains small extracellular vesicle (sEV) markers. (**A**) Cell viability analysis of RS cells 72 hours post-irradiation following a 24-hour pre-treatment with concentrated conditioned media (CCM) from RS (SF7761), RS-Clone 1, or RR (SF8628) cells. Viability was assessed after exposure to 4 Gy or 8 Gy irradiation. Data are presented as the means ± SEM. Statistical significance was determined using two-way ANOVA followed by Tukey’s post hoc test \*\**p* < 0.01, \*\*\**p* < 0.001. (**B**) Representative transmission electron microscopy (TEM) images of sEVs isolated from RR cells using differential ultracentrifugation. Red boxes indicate vesicular structures. (**C**) Particle size distribution and concentration of sEVs isolated from RS, RS-Clone 1, and RR cells, as measured by nanoparticle tracking analysis using the Zetaview system. (**D**) Western blot analysis of whole-cell lysates (WCL) and isolated sEVs from RS and RR cells. Blots were probed for the sEV markers indicated.

Components of conditioned media such as small extracellular vesicles (sEVs) can mediate oncogenic signaling in pediatric glioblastoma^12^. Transmission electron microscopy (TEM) revealed characteristic sEV morphology^28^ in CCM-derived isolates (Figure 2B), confirming the presence of vesicular structures. We next isolated and characterized sEVs from RS-SF7761, RS-Clone 1, and RR cells using differential ultracentrifugation followed by 100 kDa ultrafiltration. Nanoparticle tracking analysis (NTA) showed similar particle size distributions across the three lines, with a peak between 100-150nm (Figure 2C). Further, particle size analysis across a panel of H3K27M-pDMG lines and subclones showed consistent sEV release, indicating that sEV production is a common feature of these tumors (Figure S2). Immunoblot analysis confirmed the presence of canonical EV markers, including tetraspanins (CD9, CD81), calnexin, and histone 3 (Figures 2D and S3). Interestingly, EV marker CD63 was absent, suggesting this may be a tumor-specific feature, while CD81 was in enriched in RR-derived sEVs.

Although RS-Clone 1 is intrinsically radioresistant, its CCM did not significantly enhance survival in recipient cells, suggesting that intrinsic radioresistance does not necessarily correlate with strong paracrine signaling. In contrast, CCM from the fully RR line significantly promoted radiation survival, indicating that it may contain both EV-associated and non-EV factors to enhance recipient cell survival. To determine whether sEVs specifically contribute to these effects, we next examined whether sEVs isolated from RR cells and RS-Clone 1, independent of other secreted factors, could be internalized and influence the radiosensitivity of SF7761 cells.

### 3.3 Radiosensitive H3K27M-pDMG cells internalize radioresistant sEVs via a receptor-mediated, radiation-stress induced manner

Having observed that radioresistant (RR)-derived conditioned media enhances RS cell viability, we next examined the direct uptake of sEVs between H3K27M-pDMG cells by labeling sEVs with carboxyfluorescein diacetate succinimidyl ester (CFDA-SE), a membrane permeable dye that binds to intracellular macromolecules upon hydrolysis. Because free CFDA-SE dye can produce false positives^29^, we validated our labeling method using multiple control conditions: RS cells treated with either unlabeled sEVs (negative control), CFDA-SE-labeled sEVs retained by a 100 kDa filter (100 kDa concentrate), the 100 kDa ultrafiltration flow-through (100 kDa Filtrate), and directly stained RS cells (positive control). Flow cytometry confirmed significantly higher mean fluorescence intensity (MFI) in RS cells treated with the 100 kDa concentrate population compared to both the negative control and 100 kDa filtrate populations (p <0.001; Figure S4A-B).

To determine whether internalized sEVs localize intracellularly or remain membrane-bound, we performed confocal microscopy of RS cells stained with of DAPI and phalloidin. Imaging confirmed intracellular localization of labeled RR-sEVs (Figure 3A). High-throughput imaging of hundreds of RS cells further demonstrated a significant increase in intracellular CFDA-SE signal after RR-sEV treatment, indicating robust uptake of RR-sEVs by RS cells (p <0.001 compared to negative control; Figures 3B and S4C). To assess whether radiation stress influences sEV uptake dynamics, we irradiated RS cells (4 Gy) and exposed them to RR-sEV, with internalization measured over time. As early as 2 hours post-irradiation, sEV uptake was significantly elevated compared to non-irradiated controls (*p* <0.001; Figure 3C). By 15 hours, however, uptake levels between irradiatied and non-irradiated cells showed no significant difference, suggesting a transient but pronounced effect of radiation on early sEV internalization.

**Figure 3:**
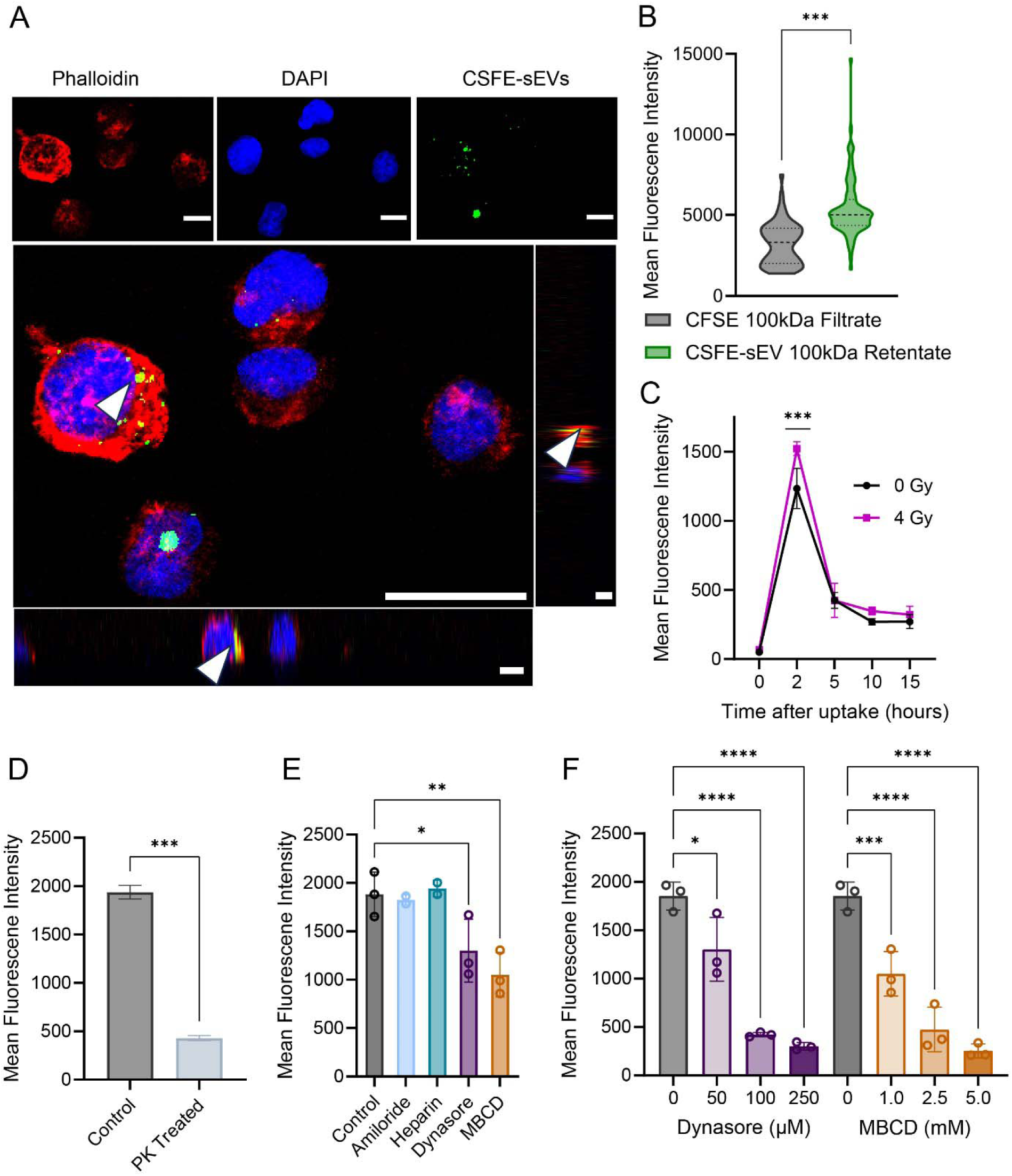
Radiosensitive H3K27M-pDMG cells internalize radioresistant sEVs through a receptor-mediated, radiation stress-induced mechanism. (**A**) Confocal microscopy images of RS cells treated with CFSE-labeled RR-sEVs for 2 hours. Cells were stained with DAPI (nuclei, blue) and phalloidin (actin, red). Adjacent orthogonal sections of a maximum Z-projection highlight intracellular vesicle localization (arrowheads). Scale bar = 10 µm. (**B**) Quantification of mean fluorescence intensity (MFI) of the CFSE signal in RS cells, comparing uptake between CSFE-100kDa filtrate (negative control) and CSFE-sEV-100kDa retentate treated cells. (**C**) Quantification of MFI measuring CSFE signal between irradiated (4 Gy) and non-irradiated (0 Gy) conditions. (**D**) MFI quantification of RS cells treated with CFSE-labeled RR-sEVs (control) or proteinase K-treated CFSE-labeled RR-sEVs (PK-treated) after 2 hours, demonstrating receptor-mediated uptake. (**E**) MFI quantification of RS cells treated with CFSE-labeled RR-sEVs (control) or pretreated for 30 minutes with uptake inhibitors: Amiloride (100 µM), Heparin (5 µg/mL), Dynasore (50 µM), or MBCD (1 mM). (**F**) MFI quantification of RS cells treated with CFSE-labeled RR-sEVs (control) or pretreated with increasing concentrations of Dynasore (0, 50, 100, 250 µM or increasing concentrations of MBCD (0, 1.0, 2.5, 5.0 mM). All experiments were performed in at least biological triplicate. Statistical significance was determined using one-way ANOVA followed by Tukey’s post hoc test. \**p* < 0.05, \*\**p* < 0.01, \*\*\**p* < 0.001, \*\*\*\**p* < 0.0001.

To determine whether sEV uptake is receptor-mediated, we pre-treated RR-sEVs with Proteinase K, which degrades surface proteins. This treatment significantly reduced sEV internalization (p <0.01; Figure 3D) indicating that sEV surface proteins are necessary for uptake. We next tested a panel of endocytosis inhibitors to determine which pathways were involved (Figure 3E). Clathrin/caveolin inhibition (dynasore) and lipid raft disruption (methyl ß-cyclodextrin, MBCD) dose-dependently blocked sEV uptake (*p* <0.05 – *p* <0.001; Figure 3F), suggesting that endocytosis through these pathways plays a major role. By contrast, amiloride and heparin had little impact, implying that macropinocytosis and heparan sulfate proteoglycan interactions are less relevant for sEV uptake in this model.

Collectively, these findings indicate that H3K27M-pDMG cells actively internalize sEVs, that radiation stress transiently enhances sEV uptake within the first few hours, and uptake primarily occurs through receptor-mediated endocytic pathways.

### 3.4 H3K27M-pDMG sEVs cargo profiling reveals glycolytic proteins, oncogenic miRNAs, and unique small molecules

sEVs carry a diverse range of cargo, including proteins, nucleic acids, and metabolites^30^. To date, only three H3K27M-pDMG cargo profile datasets exist, all of which focus soley on small RNA^12,15,16^. To expand this understanding, we profiled the proteomic and micro RNA composition of sEVs from radiosensitive (RS), partially resistant (RS-Clone 1), and fully radioresistant (RR) cell lines. Additionally, given the pronounced secretome phenotype observed in RR cells, we performed untargeted metabolomics on RR-derived sEVs (Table 1).

**Table 1.**
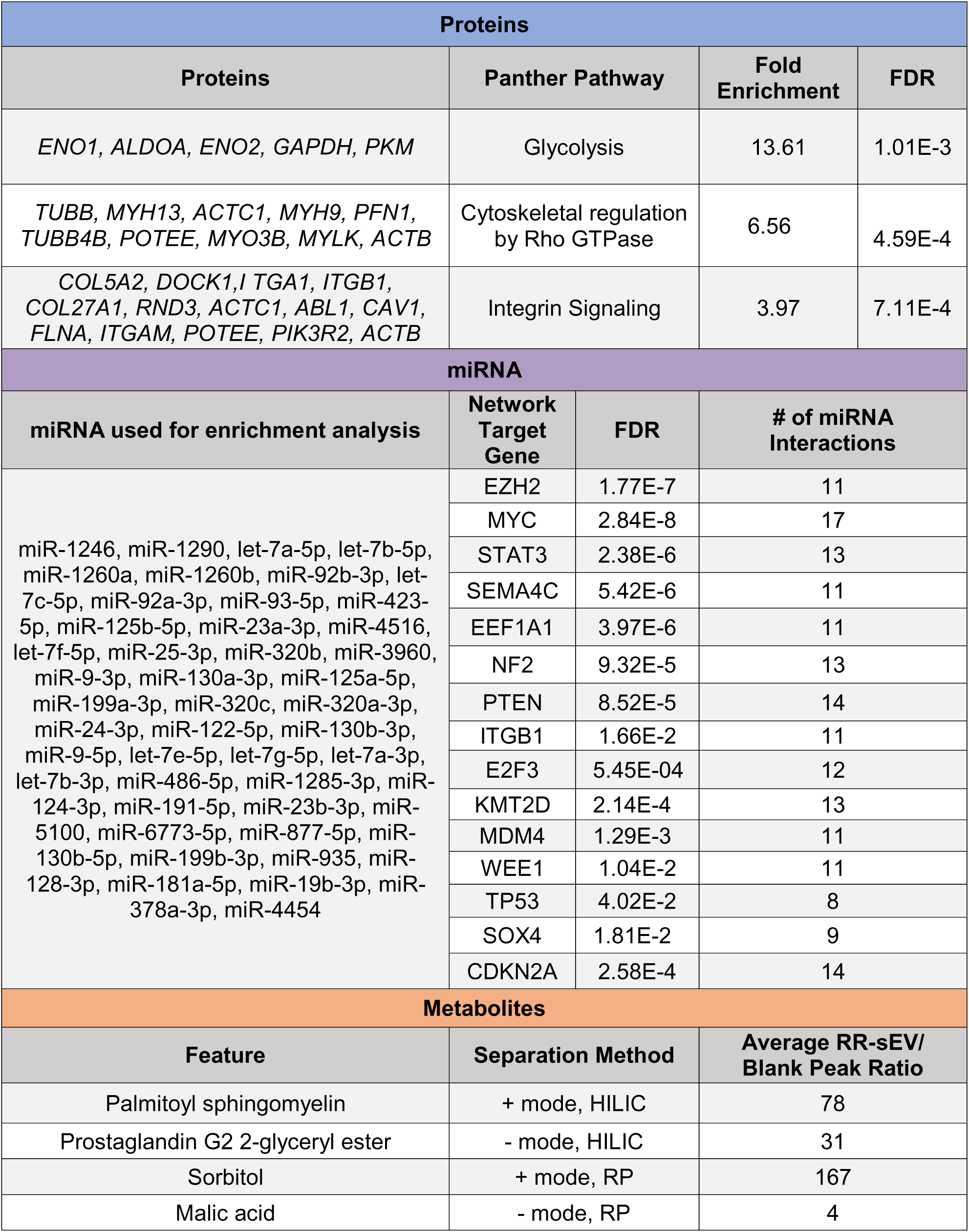
RR-sEV Enriched Proteomic, miRNA, and Metabolite Features.

LC-MS/MS identified 382 proteins in RR-sEVs, 339 in RS-Clone 1 sEVs, and 916 in RS-sEVs (Tables S1). Less than 10% of these proteins mapped to plasma membrane markers (Figure S5A), suggesting an enrichment of intracellular derived cargo. Many of these proteins were linked to caveolin-, clathrin-, and lipid raft-mediated endocytosis pathways, which were inhibited by dynasore and MBCD treatment in our uptake assays (Figure 3F). A core set of 31 proteins was shared among all three groups, and gene ontology analysis (g:Profiler) confirmed their classification as EV-related (Figure S5B). Pathway enrichment analysis (PANTHER) showed that glycolysis and integrin signaling were significantly enriched in RS-sEV and RS-Clone 1-sEV (FDR<0.05; Figures S5C-D). RR-sEVs contained a more diverse protein cargo (>20 representatives each), with significant enrichment in glycolysis, integrin signaling, and cytoskeletal regulation by Rho GTPase, suggesting broad functional capabilities that may influence recipient cells (Table 1).

Because micro RNAs (miRNAs) play a well-established role in tumor progression^31^ and sEVs serve as miRNA carriers^32^, we next characterized the miRNA content of RR-, RS- and RS-Clone 1 sEVs. Several highly expressed miRNAs were unique to RR-sEVs, while others were shared across all groups (Table S2). Notably, multiple RR-sEV miRNAs exhibited >1 logCPM (counts per million) expression after normalization. A miRNA-gene interaction network revealed that several of these miRNAs target key oncogenic pathways, including *EZH2*, *MYC*, and *TP53* (Table 1), suggesting a potential role in promoting glioma aggressiveness and therapy resistance.

To further explore the metabolic content of H3K27M-pDMG sEVs, we performed untargeted metabolomics on RR-sEVs using four LCMS/MS methods spanning both polar and non-polar metabolites^33^. Across reverse-phase positive (RP-Pos), reverse-phase negative (RP-Neg), hydrophobic interaction liquid chromatography negative (HILIC-Neg), and a HILIC positive (HILIC-Pos) ion modes, we detected 832, 244, 409, and 1057 unique molecular features, respectively (Table S3). Several metabolites exhibited fold changes significantly higher than background controls (Table 1). Among these, palmitoyl sphingomyelin, a C16 ceramide, was enriched in RR-sEVs. Alterations in ceramide equilibrium have been implicated in cancer progression, with shifts towards sphingomyelin species promoting tumor survival^34^.

Together, these findings demonstrate that H3K27M-pDMG-derived sEVs carry a diverse array of biologically relevant cargo, including glycolytic and integrin-associated proteins, oncogenic miRNAs, and bioactive metabolites. These data highlight both intra- and intertumoral variation in sEV composition and suggest sEV cargo may play an active role in H3K27M-pDMG pathophysiology.

### 3.5 Radioresistant sEVs induce transient metabolic changes and long-term transcriptional reprogramming in radiosensitive H3K27M-pDMG cells

Since our uptake studies demonstrated robust sEV internalization within 24 hours (Figure 3) and cargo profiling identified enrichment in oncogenic signaling pathways (Table 1), we next examined whether RR-sEVs alter the metabolic state of recipient RS cells. Cells were treated with RR-sEVs under the same conditions as uptake assays, and intracellular metabolites were extracted 4 hours post-treatment (Figure 4A). Partial least squares discriminant analysis (PLS-DA) showed clear metabolic separation between RR-sEV and control RS cells, indicating that RR-sEVs shift the intracellular metabolite profile after RR-sEV exposure (Figure 4B). Hierarchical clustering showed broad alterations in metabolite abundance, including amino acids, sugars, and lipid intermediates (Figure 4C). Subsequent pathway enrichment analysis highlighted upregulation of processes related to glycolysis, mitochondrial shuttles, and lipid metabolism (Figure 4D). Notably, these metabolic differences were transient and not sustained at 16 hours post-treatment (data not shown), suggesting that while a single treatment with RR-sEVs can trigger early metabolic fluctuations, they do not lead to long-term metabolic reprogramming in this experimental setting. However, the acute nature of these changes raised the possibility that they might precede or prime longer-term transcriptional effects.

**Figure 4:**
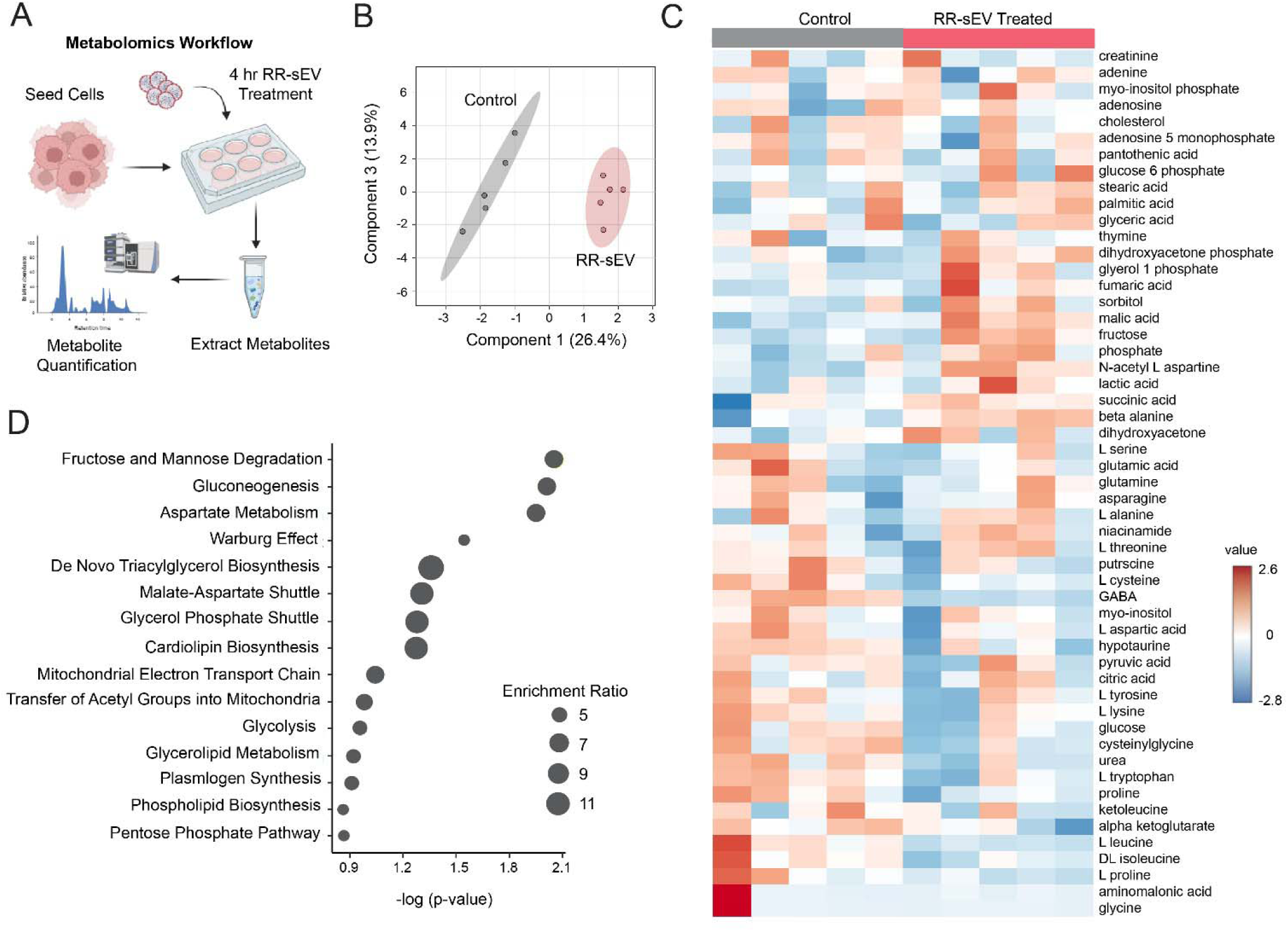
RR-sEVs induce metabolic remodeling in radiosensitive cells. (**A**) Schematic representation of the metabolomics workflow. RS cells were treated with RR-sEVs for 4 hours prior to metabolite extraction and analysis. (**B**) Partial Least Squares Discriminant Analysis (PLS-DA) of global metabolite profiles showing distinct clustering of RR-sEV treated RS cells compared to control (n = 5 per group). (**C**) Heatmap of standardized metabolite abundance (Z-scores) showing relative levels of detected metabolites in RR-sEV treated and control cells. (**D**) Pathway enrichment analysis of differentially regulated metabolites in RR-sEV treated RS cells compared to control. Size of circles reflects enrichment ratio, and position on the x-axis reflects statistical significance (FDR <0.25) All metabolite concentrations were normalized to internal standards and using one-factor statistical analysis in MetaboAnalyst. Control samples received the EV-depleted 100 kDa filtrate fraction.

To investigate this, we performed RNA-seq at 18 hours post-treatment (Figure 5A). Principle component analysis showed distinct clustering of RR-sEV-treated cells away from RS-Clone 1-sEV and control groups, suggesting broad transcriptional shifts (Figure 5B). Differential expression analysis identified 52 significantly regulated genes (FDR < 0.05) in RR-sEV treated cells, with notable upregulation of genes encoding NADH:ubiquinone oxidoreducatase (Complex I) subunits (Figure 5C), which are central to mitochondrial oxidative phosphorylation and have been linked to therapy resistance. Gene set enrichment analysis (GSEA) further supported this observation, identifying oxidative phosphorylation as a top enriched pathway in RR-sEV treated cells (Figure 5D; NES: 1.58, FDR: 0.05). The enrichment pattern, driven largely by mitochondrial and electron transport chain-associated genes, aligns with early metabolomic findings suggesting mitochondrial involvement.

**Figure 5:**
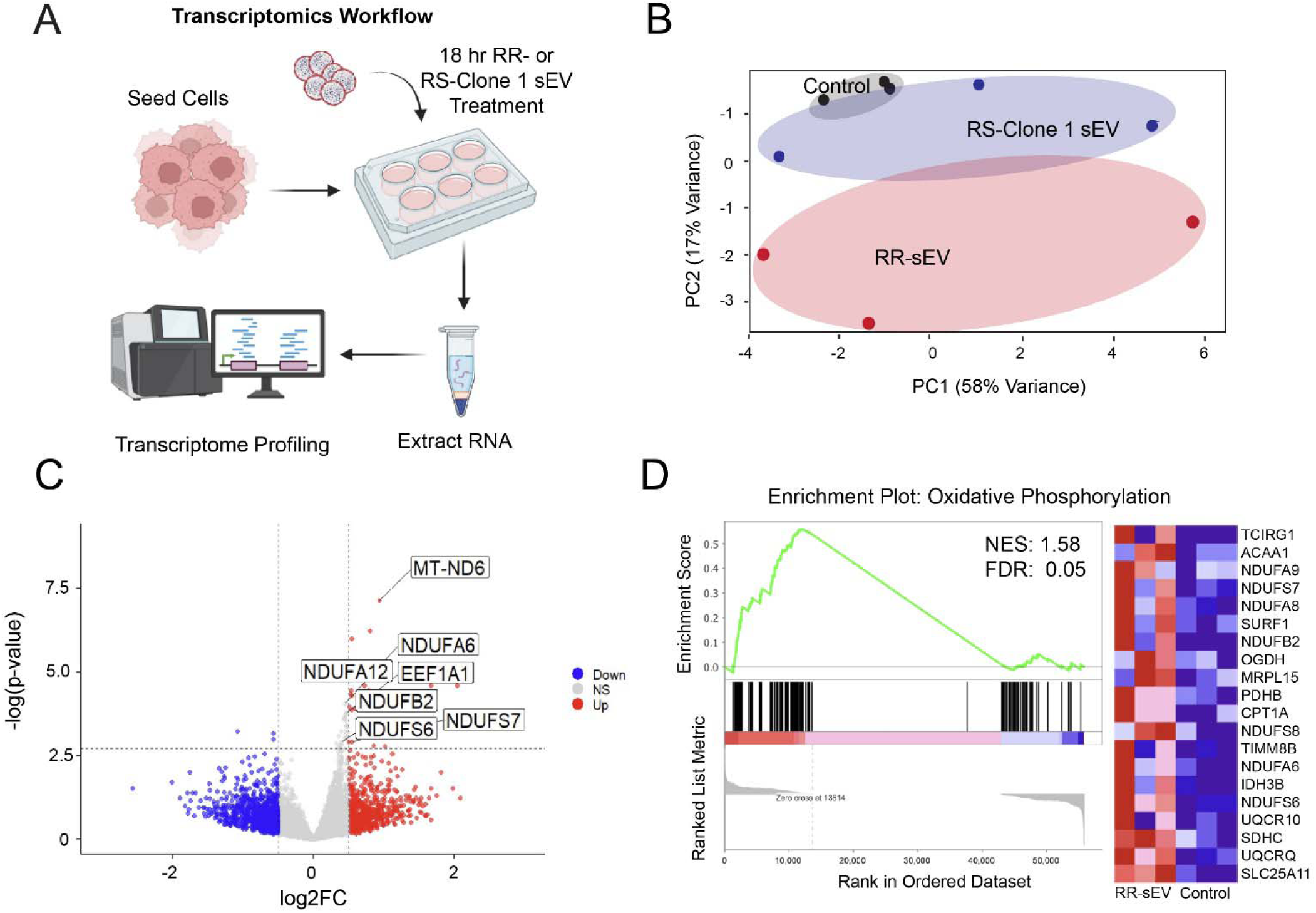
RR-sEVs induce transcriptional changes linked to mitochondrial metabolism in radiosensitive cells. (**A**) Schematic representation of the transcriptomic workflow: RS cells were treated with RR-sEVs, RS-Clone 1-sEVs, or control (EV-depleted 100 kDa filtrate) for 18 hours prior to RNA extraction and bulk RNA sequencing. (**B**) Principal component analysis (PCA) of transcriptomes shows distinct clustering of RR-sEV-treated RS cells compared to RS-Clone 1-sEV and control groups, indicating broad transcriptional shifts (n = 3 per group). (**C**) Volcano plot displaying differentially expressed genes in RR-sEV-treated cells versus control. Red points indicate significantly upregulated genes and blue points indicate downregulated genes (FDR < 0.05; vertical dashed lines denote log2 fold change thresholds of ±1). (**D**) Gene set enrichment analysis (GSEA) identified enrichment of oxidative phosphorylation gene signatures in RR-sEV-treated cells (NES = 1.58, FDR = 0.05), including increased expression of mitochondrial complex I subunit genes. Heatmap at right displays top 20 genes contributing to enrichment.

Interestingly, sEVs derived from the partially radioresistant subclone RS-Clone 1 did not induce comparable transcriptional changes (Figure 5B), highlighting the complexity and specificity of sEV-mediated effects on recipient cell phenotypes. These findings suggest that RR-sEVs activate a mitochondrial transcriptional program that may enhance energy production and stress resilience. While a single RR-sEV exposure may not be sufficient to drive lasting metabolic reprogramming, repeated or sustained exposure *in vivo* could contribute to metabolic plasticity and the emergence of therapy-resistant phenotypes within heterogenous tumor microenvironments.

### 3.6 Radioresistant sEVs enhance DNA damage resolution in radiosensitive cells

Our previous findings demonstrated that RR-sEVs induce transcriptional shifts in recipient RS cells, particularly in pathways related to oxidative phosphorylation. Given that mitochondrial metabolism is intricately linked to the DNA damage response (DDR)^35^, we next investigated whether RR-sEVs modulate the cellular response to ionizing radiation. Specifically, we assessed the accumulation and repair of radiation induced double-strand breaks (DSBs) using two canonical markers, γ-H2AX and 53BP1^36^. As shown in Figure 6A, γ-H2AX serves as a sensor of DNA damage, rapidly accumulating at DSB sites and facilitating the recruitment of repair machinery^39,40^. 53BP1 is a core mediator of the repair process itself, forming puncta at DSB sites and promoting non-homologous end joining to maintain genome stability^37,38^.

**Figure 6:**
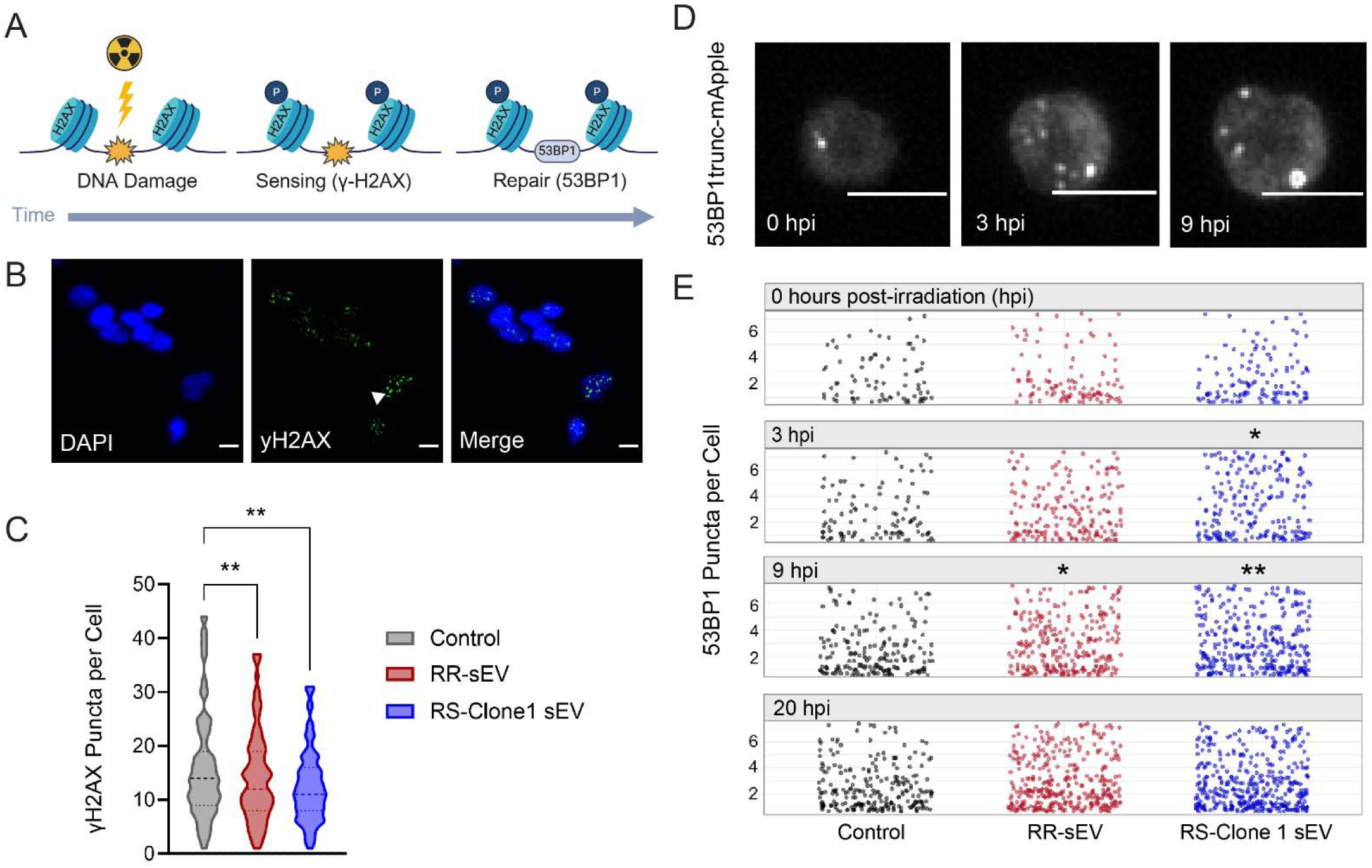
RR-sEVs enhance the DNA damage response in radiosensitive H3K27M-pDMG cells. (**A**) Schematic representation of radiation-induced DNA damage and repair over time, highlighting γ-H2AX formation as a DNA damage sensor that promotes 53BP1 recruitment to facilitate repair. (**B**) Representative immunofluorescence images of RS cells stained for DAPI (blue) and γ-H2AX (green) 4 hours after 4 Gy irradiation, following 18-hour pre-treatment with either control, RR-sEVs, or RS-Clone 1-sEVs. Arrowhead indicates γ-H2AX foci. Scale bar = 10 μm. (**C**) Quantification of γ-H2AX puncta per cell across the three treatment groups. (**D**) Time-lapse imaging of 53BP1trunc-mApple puncta in RS cells before and after irradiation (0, 3, and 9 hours post-irradiation). Scale bar = 10 μm. (**E**) Quantification of 53BP1 puncta per cell over time (0, 3, 9, and 20 hours post-irradiation) in RS cells pre-treated with control, RR-sEVs, or RS-Clone 1-sEVs. All experiments were conducted with a starting cell density of 2,500 cells per well, with at least two technical replicates and three independent biological replicates. The control condition consisted of the EV-depleted 100 kDa filtrate fraction. Data in C are presented as mean ± SEM. Statistical analysis for γ-H2AX puncta was performed using one-way ANOVA with Tukey’s post-hoc test. Statistical analysis for 53BP1 puncta (E) was performed using two-way ANOVA followed by Tukey’s post-hoc test. \**p* = 0.05, \*\**p* < 0.01, compared to control.

To quantify DNA damage, we first identified 4 hours post-irradiation (hpi) as the peak of γ-H2AX accumulation in RS cells and used this timepoint for comparative analysis. RS cells pre-treated with RR-sEVs or RS-Clone 1-sEVs exhibited significantly fewer γ-H2AX foci per cells compared to control-treated cells (*p* < 0.01) after 4Gy irradiation, suggesting a reduction in persistent DSBs (Figure 6B-C). We next evaluated the kinetics of DNA repair by tracking 53BP1 puncta in live RS cells stably expressing a fluorescent 53BP1trunc-mApple reporter. Cells were treated with RR-sEVs, RS-Clone 1-sEVs, or vehicle control for 18 hours, irradiated with 4Gy, and imaged at 0, 3, 8, and 20 hpi (Figure 6D-E). At baseline (0 hpi), no differences in puncta formation were observed among groups. However, by 3 hpi, RS-Clone 1-sEV treated cells exhibited significantly more 53BP1 puncta per cell than controls (*p* = 0.03). At 9 hpi, both RR-sEV and RS-Clone 1-sEV treated cells had significantly more puncta than controls (*p* = 0.03, *p* = 0.003, respectively). By 20 hpi, the number of 53BP1 puncta per cell had returned to baseline levels across the groups.

Together, these findings suggest that sEVs from both RR and RS-Clone 1 cells promote DDR capacity in recipient radiosensitive cells by both reducing persistent DNA damage, as evidenced by lower γ-H2AX levels, and promoting more robust or sustained repair activity, as indicated by elevated 53BP1 puncta. These findings support a model in which sEVs derived from radioresistant subpopulations contribute to adaptive resilience within heterogenous tumor microenvironments by facilitating intercellular transfer of repair-promoting signals.

### 3.7 Radioresistant sEVs delay radiation-induced death in radiosensitive cells

Since RR-sEVs influence metabolism, transcriptional programs, and DNA damage resolution, we next examined their functional significance on radiosensitive (RS) SF7761 cells during acute radiation exposure. Given their enrichment of oxidative phosphorylation and DNA repair pathways in recipient cells, we hypothesized that RR-sEVs may enhance cell survival under radiation stress. Before assessing sEV-mediated effects on radiation-induced cell death, we first evaluated whether bulk cell viability assays could accurately capture sEV-specific effects. We tested sEV suspensions without cells in two widely used assays: Caspase-Glo, which detects caspase 3/7 activity as a marker of apoptosis and CellTiter-Glo, which measures cellular ATP as an indicator of viability. sEV suspensions generated signals in both assays, with RR-sEVs eliciting a robust ATP signal (Figure S6A) and a dose-dependent increase in caspase activity, surpassing control and conditioned media signals (Figure S6B). These findings indicate that sEVs themselves can contribute to assay signals, potentially confounding the interpretation of traditional cell population-based viability assays.

To overcome the limitations and directly measure single cell responses, we utilized a Genetically Encoded Death Indicator (GEDI)^41^, a dual-fluorescence biosensor that tracks apoptotic commitment based on fluorescence ratio shifts. The GEDI construct contains both a genetically encoded calcium sensor (GC150) and the red fluorescent protein mApple, which serves as a morphological marker. Apoptotic commitment is indicated by an irreversible loss of membrane homeostasis, leading to an increase in intracellular calcium levels. This results in an enhanced GC150 fluorescence signal, which, when normalized to mApple, generates a GEDI ratio that reliably predicts cell fate (Figure 7A). We empirically determined a GEDI threshold corresponding to cell death by using an extreme radiation dose (25 Gy). Cells with a GEDI ratio of >1 were classified as apoptotic, while those maintaining a ratio of <1 were considered viable (Figure 7B-D). This single-cell approach provides a high-resolution, quantitative readout of radiation-induced cytotoxicity, independent of confounding factors introduced by sEVs in bulk assays.

**Figure 7:**
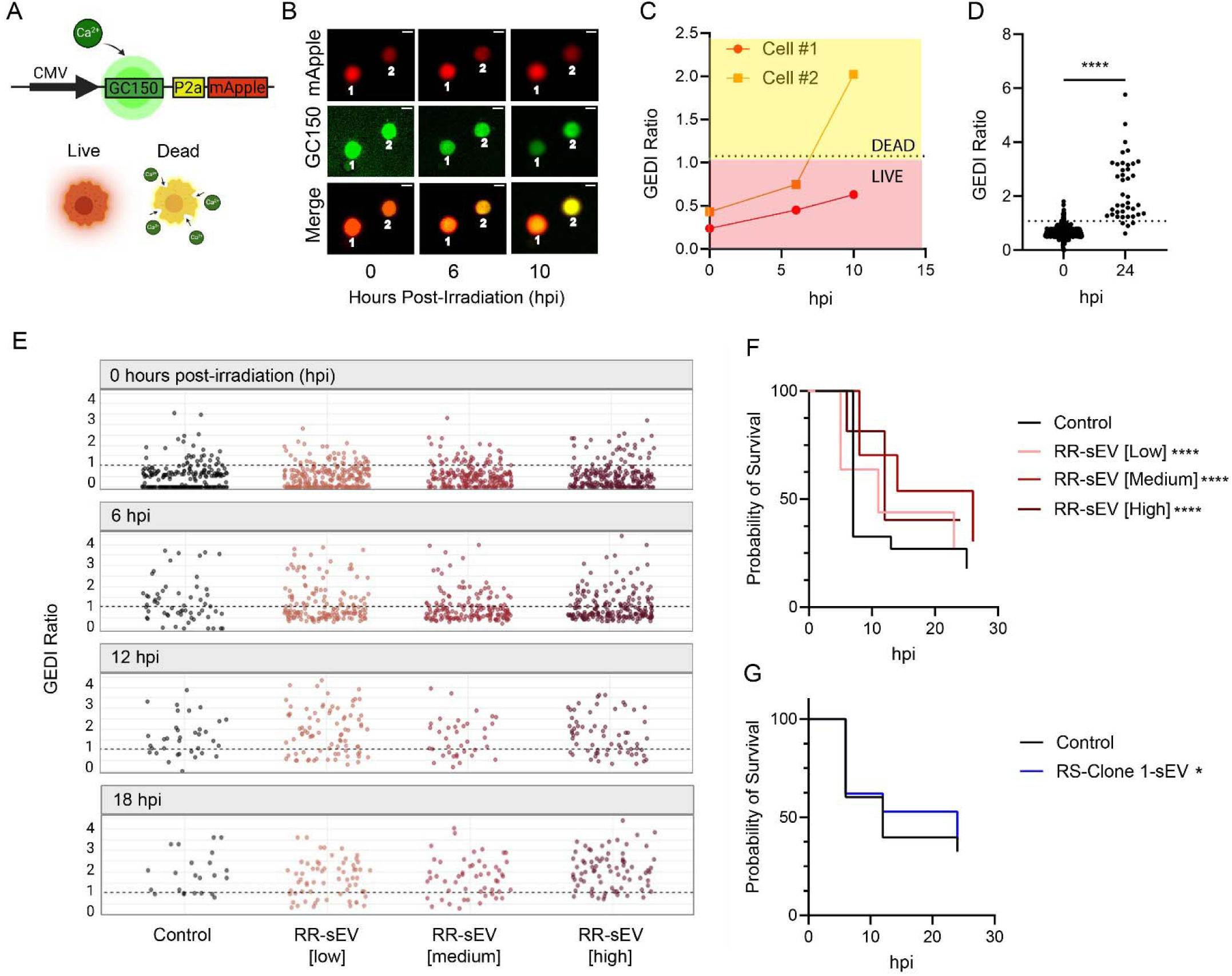
RR-sEVs delay radiation-induced cell death in radiosensitive cells. (**A**) Schematic of the Genetically Encoded Death Indicator (GEDI) construct used in this study. Upon lethal intracellular calcium influx, GC150 fluorescence increases, resulting in a shift in the GEDI ratio (GC150/mApple) that distinguishes live from dying cells. (**B**) Representative time-lapse images of RS cells stably expressing GEDI, showing a transition from live (low GEDI ratio) to dead (high GEDI ratio) status following exposure to 8 Gy radiation. Scale bar = 10 μm. (**C**) GEDI ratio plotted for two individual cells from (B), illustrating dynamic shifts in intracellular calcium over time. (**D**) GEDI ratio of individual RS cells 24 hours after 25 Gy irradiation, compared to baseline (0 hpi). Dotted line indicates the empirically defined GEDI threshold. (**E**) GEDI ratio values from single RS cells pretreated with control or increasing doses of RR-sEVs (low, medium, high), measured at 0, 6, 12, and 18 hours post-irradiation (hpi). The progressive reduction in datapoints over time reflects cell death and subsequent detachment from the plate. (**F**) Kaplan-Meier survival analysis of RS-GEDI cells treated with three different doses of RR-sEVs or control prior to irradiation. (**G**) Kaplan-Meier survival analysis of RS-GEDI cells treated with RS-Clone 1-sEVs or control prior to irradiation. All experiments were performed in biological triplicate with ≥2 technical replicates per condition, using a seeding density of 2,500 cells per well. Statistical analysis in (D) was performed using a paired t-test. Statistical analysis in (F–G) was performed using the log-rank test. Significance is indicated as \**p* = 0.05, \*\*\**p* < 0.001, \*\*\*\**p* < 0.00001 compared to control values.

To determine whether RR-sEVs influence radiosensitivity, RS cells were treated with three doses of RR-sEVs (low: 2,500 sEVs/cell, medium: 5,000 sEVs/cell, high:15,000 sEVs/cell) for 24 hours prior to irradiation. Cell fate was monitored at 0, 6, 12, and 18 hours post-irradiation. In untreated cells, the GEDI ratio increased over time, reflecting progressive radiation-induced cell death, whereas RR-sEV treated cells maintained lower GEDI ratios (e.g. at 6 hpi: untreated = 1.76, low-dose = 1.18, *p* < 0.001, high-dose = 1.2, *p* < 0.001) (Figure 7E). A Kaplan-Meier survival analysis was conducted based on single-cell outcomes, with cells classified as alive (GEDI ratio < 1) or dead (disappearing from the field of view). Log-rank testing confirmed a significant increase in survival probability in RR-sEV-treated cells in a dose-dependent manner (*p* < 0.001; Figure 7F). To determine whether this effect was reproducible with clonal sEVs, RS cells were treated with RS-Clone 1 sEVs and analyzed in the same manner (Figure S7). In contrast to the pronounced effects of RR-sEVs, RS-Clone 1 sEVs conferred a modest survival advantage (*p* = 0.05; Figure 7G).

These results highlight the limitations of traditional bulk cell population assays in detecting sEV-specific effects and demonstrate the value of biosensors such as GEDI as a high-resolution tool for tracking radiation-induced cytotoxicity. More importantly, these findings provide direct evidence that RR-sEVs promote radioresistance of RS cells in a dose dependent manner, consistent with their observed ability to enhance oxidative phosphorylation and accelerate DNA repair. In contrast, the more modest effect of RS-Clone 1 sEVs suggests that not all tumor-derived sEV populations exert equal protective effects, emphasizing the heterogeneity of sEV-mediated paracrine signaling within the tumor microenvironment.

## 4 Discussion

In this study, we investigated whether small extracellular vesicles derived from radioresistant (RR) H3K27M-pDMG cells can confer radioprotective effects to a radiosensitive population. Using a treatment-naïve H3K27M-pDMG model, we demonstrated that RR-sEVs promote multiple pro-survival phenotypes under radiation stress. Specifically, RR-sEV treatment rapidly shifted TCA-cycle metabolism, upregulated gene sets involved in oxidative phosphorylation and MYC pathways, and enhanced DNA damage resolution via increased 53BP1 puncta and decreased γ-H2AX foci. This ultimately led to delayed radiation-induced cell death at the single-cell level. Although sEVs from a partially resistant subclone (RS-Clone 1) of the treatment-naïve, radiosensitive line exhibited some similar effects, they lacked the pronounced protective influence observed with RR-sEVs, indicating variability in sEV-mediated signaling among intratumoral populations.

Our findings align with previous reports that implicate sEVs in oncogenic reprogramming and therapy resistance in other tumor types, including adult gliomas, prostate cancer, and oral squamous carcinoma. sEVs can facilitate tumor progression by delivering oncogenic proteins such as EGFR and ANXA2 to promote angiogenesis^42,43^, activate intracellular signaling pathways such as the PI3/Akt axis to enhance DNA repair and cell proliferation^44^, and suppress apoptosis through microRNA-mediated regulation^45^. In pediatric high-grade gliomas, prior studies have shown that H3K27M-pDMG-derived sEVs influence gene expression in neural stem cells^15^ and exhibit subclone-specific miRNA content^12^. However, the direct impact of H3K27M-pDMG-derived sEVs on tumor radioresistance has remained largely unexplored. Our study addresses this gap by demonstrating that RR-sEVs are internalized via receptor-mediated endocytosis, particularly under radiation stress, and that their cargo, including oncogenic miRNAs, glycolytic proteins, and lipid species, can reprogram recipient cells towards a more radioresistant phenotype.

The ability of RR-sEVs to enhance radiation resistance appears to be mediated, in part, through metabolic reprogramming, we observed shifts in TCA cycle metabolites at early time points, with subsequent transcriptional enrichment of oxidative phosphorylation pathways, suggesting a coordinated shift to mitochondrial metabolism. Increased oxidative phosphorylation has been implicated in radiation resistance across multiple tumor types, as it supplies ATP for DNA repair and mitigates oxidative damage by maintaining redox balance^46,47^. The enrichment of MYC target genes further supports role for RR-sEVs in reprogramming recipient cells, as MYC has been shown to promote metabolic flexibility and cell survival under stress conditions^48–50^. The overrepresentation of glycolytic and integrin-related cargo in RR-sEVs suggests additional roles in energy metabolism and altered adhesion dynamics, which could further support survival under radiation-induced stress^51–53^. These findings indicate that metabolic rewiring may be a key mechanism through which RR-sEVs enhances therapy resistance in recipient cells.

Beyond metabolic adaptations, RR-sEVs also influenced DNA repair processes, as evidenced by increased 53BP1 recruitment and reduced γ-H2AX foci following radiation exposure. sEVs have been shown to activate DNA repair pathways in therapy resistant models such as prostate and renal cell carcinoma by downregulating MLH1 and p53 expression^54,55^. Furthermore, we saw enrichment of gene sets related to cell differentiation, suggesting that RR-sEVs may also influence tumor plasticity, which is consistent with recent studies indicating the presence of a dynamic stem cell compartment in H3K27M-pDMG^9^. Prolonged cell cycle progression and enhanced DNA repair makes cancer stem-like cells more adept at repairing DNA damage and exhibit intrinsic resistance to therapy^56–58^. The ability of RR-sEVs to promote a more stem-like state could further contribute to tumor persistence and recurrence following radiation treatment.

Interestingly, while RS-Clone 1 sEVs share some features of RR-sEVs, including increased DNA damage resolution and over-representation of glycolytic and integrin associated proteins, they did not confer the same degree of radioprotection. Certain miRNA enriched in RR-sEVs, such as hsa-miR-1246 and hsa-miR-93-5p have known roles in promoting stemness and survival in glioblastoma^59,60^. Our analysis revealed miRNA species unique to RS-Clone 1 but not present in RR- or parental-line derived EVs, suggesting that subclonal differences in secretome profiles influence differential signaling and functional outcomes. These findings reinforce the idea that intratumoral heterogeneity in H3K27M-pDMGs is not only driven by genomic and transcriptional differences, but also by distinct secretome profiles that shape the tumor microenvironment.

The potential for sEVs to mediate therapy resistance raises important clinical considerations. Targeting sEV biogenesis, for example, through Rab27a inhibitors^61,62^ or GW4869^63^ could reduce the transfer of pro-survival signals between tumor cells.

Blocking sEV uptake, particularly through receptor-mediated endocytosis, may provide another avenue to disrupt communication between resistant and sensitive populations. Given the metabolic shifts induced by RR-sEVs, targeting oxidative phosphorylation or MYC-driven pathways could be an effective strategy for sensitizing tumor cells to radiation. Furthermore, if specific miRNAs or proteins in RR-sEVs correlate with radiation resistance, they could serve as predictive biomarkers for patient stratification, aiding in the development of more personalized therapeutic approaches.

While our study provides key insights, several limitations warrant consideration. First, although our sEV preparations exhibited low protein contamination, we did not purify them further using methods like size exclusion chromatography or density ultracentrifugation, leaving open the possibility that contaminating factors or distinct EV subpopulations may contribute to the observed effects. Future studies should characterize specific sEV subtypes to identify the most functionally relevant cargo. Second, our data indicate that sEV uptake occurs primarily via receptor-mediated mechanisms, but residual uptake following proteinase K treatment suggests alternative pathways, such as macropinocytosis, may also contribute to sEV internalization. Further investigation into the specific receptors and alternative uptake pathways will help refine our understanding of how sEVs interact with recipient cells. Lastly, our findings are limited to *in vitro* models and do not account for the complex interactions present *in vivo*, where immune cells, stromal components, and systemic factors could further influence sEV function^64^. Future work using patient-derived xenografts or organoid co-cultures will be important for validating these mechanisms in a physiologically relevant setting.

Taken together, our findings advance the understanding of sEV-mediated radioresistance in H3K27M-pDMGs. We demonstrate that sEVs derived from radioresistant tumor cells facilitate intercellular communication with radiosensitive cells, enhancing metabolic adaptability, DNA repair, and survival under radiation stress. The differential effects of RR-sEVs compared to RS-Clone 1 sEVs highlight the role of intratumoral heterogeneity therapy resistance and suggest that the ability of sEVs to promote survival is dependent on the functional properties of their cargo. These results

support the idea that targeting sEV-mediated signaling may provide a novel strategy for mediating radiation resistance in H3K27M-pDMGs. Further investigations into the specific sEV cargo components and their functional contributions to therapy resistance will be essential for developing targeted interventions that could improve outcomes for patients with this aggressive pediatric glioma.

## Author Contributions

**Viral Oza**: conceptualization (lead) data curation (lead), formal analysis (lead), investigation (lead), methodology (lead), project administration (lead), writing-original draft (lead), writing-review and editing (lead). **Yelena Chernyavskaya**: data curation (supporting), formal analysis (supporting). **Kenan A Flores** data curation (supporting), formal analysis (supporting). **Majd A Al-Hamaly** data curation (supporting), formal analysis (supporting). **Caitlyn B Smith** conceptualization (supporting), data curation (supporting), formal analysis (supporting), methodology (supporting). **Ronald C Bruntz** data curation (supporting), formal analysis (supporting), methodology (supporting), resources (supporting), software (supporting), validation (supporting). **Jessica Blackburn** conceptualization (lead), data curation (lead), formal analysis (lead), investigation (lead), methodology (lead), project administration (lead), writing-original draft (lead), writing-review and editing (lead).

## Acknowledgements

The authors acknowledge the University of Kentucky Light Microscopy Core, Bioelectronics and Nanomedicine Center, the Redox Metabolism Shared Resource, and the Flow Cytometry and Immune Monitoring core for the use of their facilities. We thank Steven Finkbeiner (The Gladstone Institutes) for the GEDI plasmid, as well as Michelle Monje (Stanford Medicine) and Nicholas Vitanza (Seattle Children’s Hospital) for H3K27M-pDMG patient cell lines.

## Data Availability

The data discussed in the present publication have been deposited in NCBI’s Gene Expression Omnibus and are accessible through GEO Series accession numbers GSE286943 (Small RNA seq), GSE286945 (Bulk Cell Line RNA Seq), and GSE286941 (EVTreat-RNASeq). All other data files and/or code supporting this study are available on reasonable request from the corresponding author.

