## Supplementary Data for "Small Extracellular Vesicles from Radioresistant H3K27M-Pediatric Diffuse Midline Glioma Cells Modulate Tumor Phenotypes and Radiation Response"

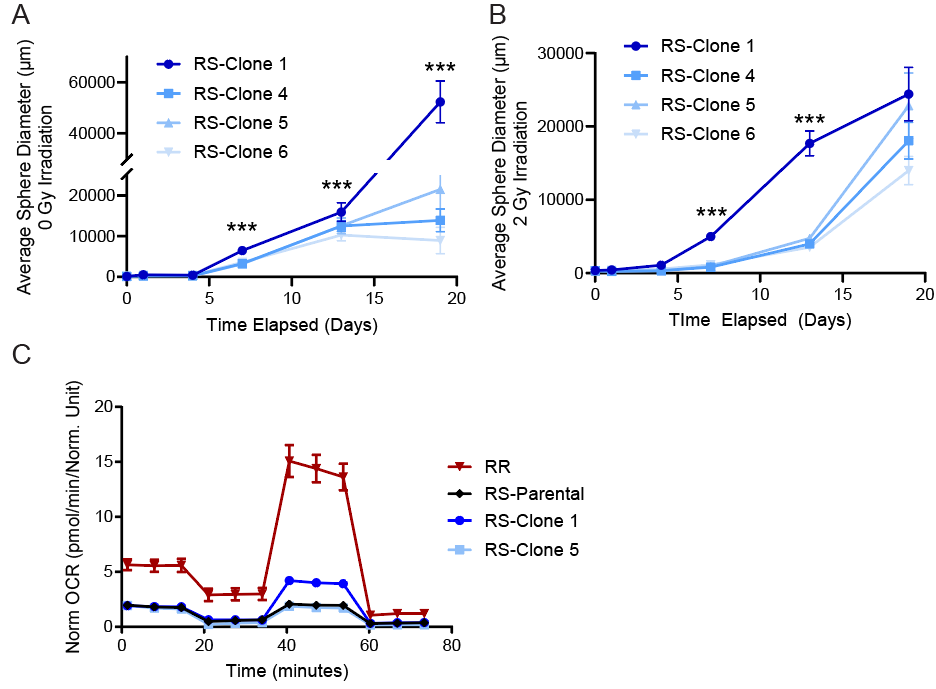

**Supplemental Figure 1: Growth and metabolic characteristics of RS-associated subclones.** (**A**) Colony formation assay showing the growth dynamics of RS-associated subclones under non-irradiated (0 Gy) conditions. Average sphere diameter was measured over time, n=3. (**B**) Colony formation assay of the same RS-associated subclones following 2 Gy irradiation, showing differential growth responses, n=3. (**C**) Mitochondrial stress test of RS-Parental, RR, and two RS-derived subclones using the Seahorse platform, n = 8. Data are presented as means ± SEM. Statistical significance was determined using two-way ANOVA followed by Tukey’s post-hoc test (**p* < 0.05, ***p* < 0.01, ****p* <0.001). All experiments were performed in biological triplicate.

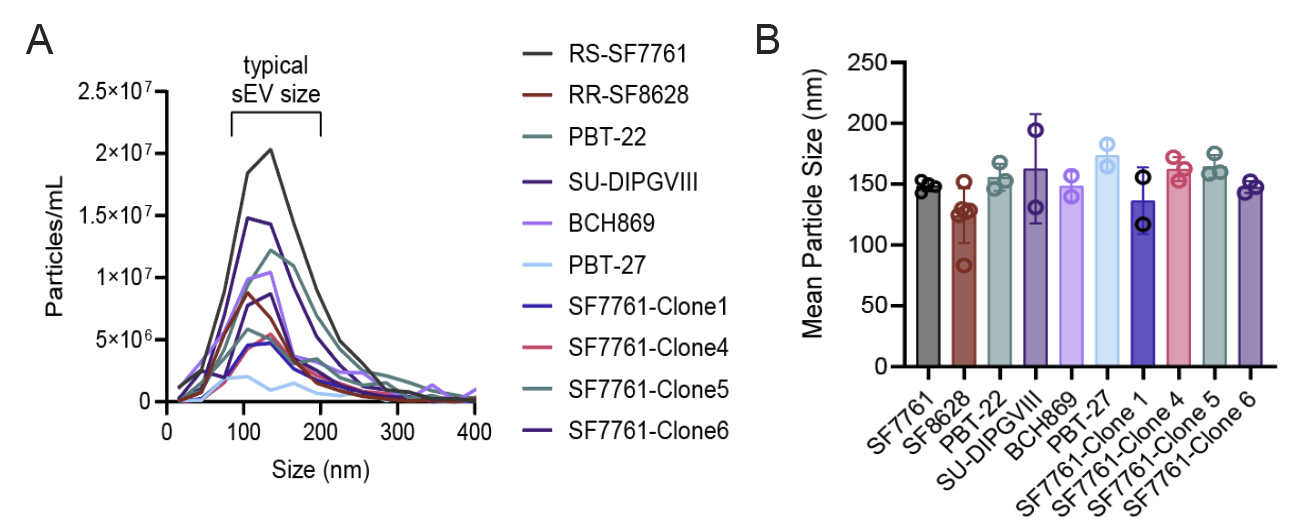

**Supplemental Figure 2: Characterization of small extracellular vesicles (sEVs) derived from H3K27M-pDMG cell lines and subclones.** (**A**) Nanoparticle tracking analysis (NTA) of sEVs isolated from multiple H3K27M-pDMG cell lines, including RS (SF7761), RR (SF8628), PBT-22, SU-DIPGVIII, BCH869, PBT-27, and four RS-derived subclones (SF7761-Clone 1, Clone 4, Clone 5, and Clone 6). The distribution of particle sizes is shown, with the typical sEV size range indicated. (**B**) Mean particle size of sEVs from each cell line, measured using the Zetaview system. Bars represent means ± standard error of the mean (SEM).

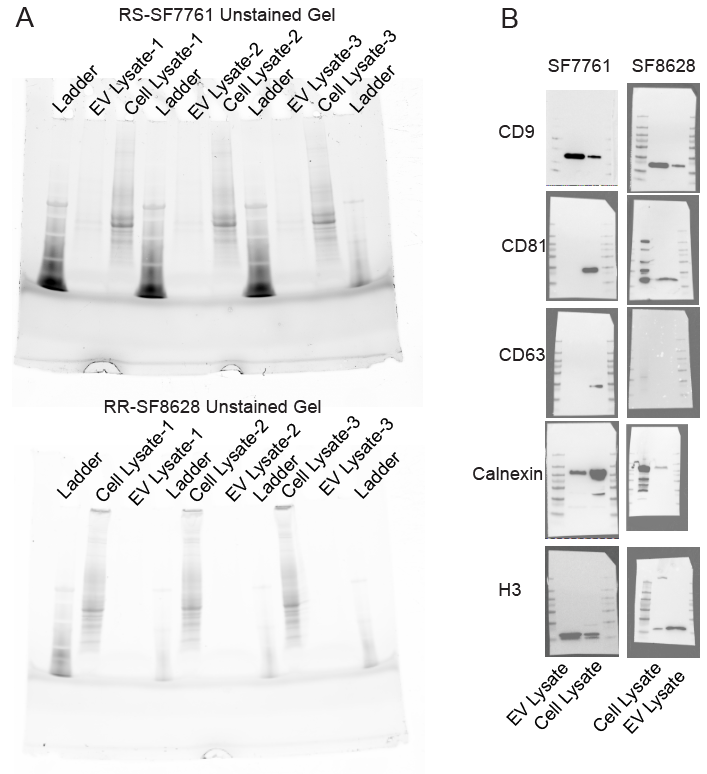

**Supplemental Figure 3: Total protein loading and blots for extracellular vesicle (EV) marker validation.** (**A**) Stain-free SDS-PAGE gel showing total protein loading of whole-cell lysates and extracellular vesicle (EV) lysates from RS (SF7761) and RR (SF8628) cells across three independent preparations. (**B**) Images of the blots corresponding to the cropped images in Figure 2, showing immunoblot detection of the indicated EV markers in whole-cell lysates (left lane) and EV fractions (right lane) from SF7761 and SF8628 cells.

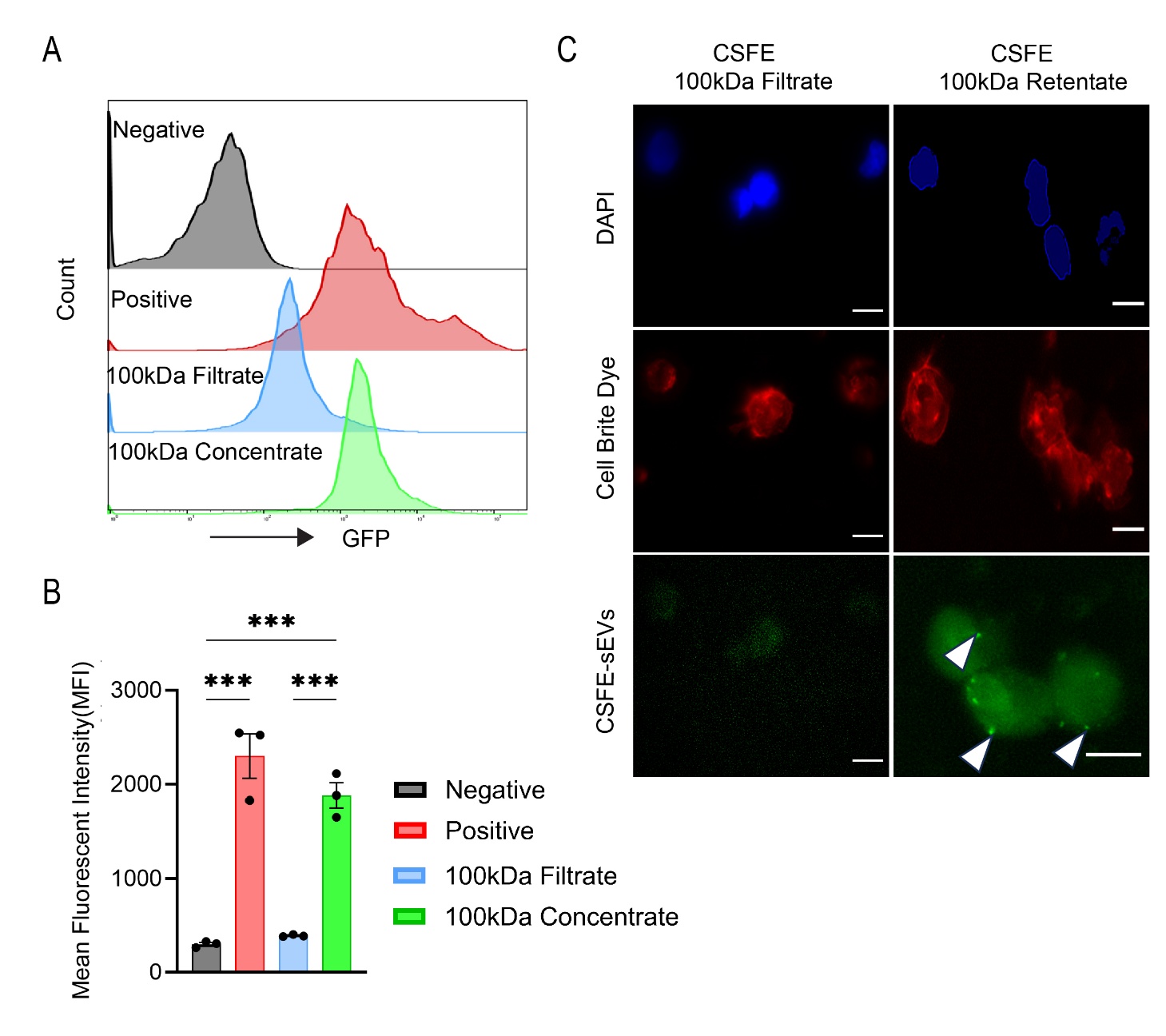
**Supplemental Figure 4: Total protein loading and blots for extracellular vesicle (EV) marker validation.** (**A**) Flow Cytometry profiles of 4 populations of RS cells, untreated (negative), cells stained with CSFE (positive), cells treated with 100 kDa filtrate of RR-sEVs dyed with CSFE (100 kDa filtrate), cells treated with 100 kDa concentrate of RR-sEVs dyed with CSFE (100 kDa concentrate). (**B**) MFI quantification of flow cytometry profiles in (A). (**C**) Representative images of high throughput immunofluorescent imaging of RS cells treated with RR-sEVs-CSFE stained with DAPI and Cell Brite. Statistical analysis performed in (B) was a 1-way Anova followed by a Tukey post-hoc. ***p<.0001. All experiments were performed in biological triplicate.

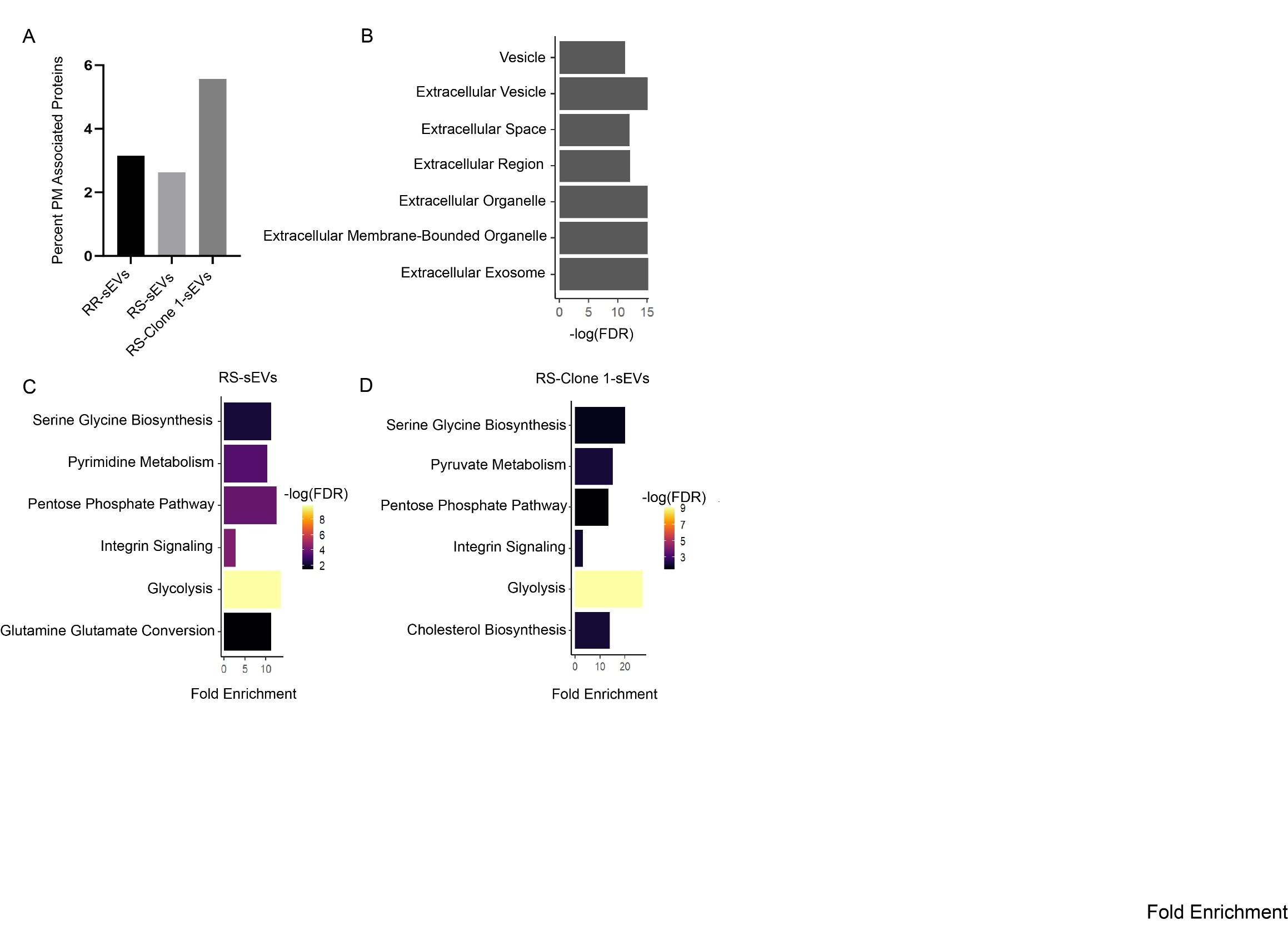

**Supplemental Figure 5: Proteomic pathway analysis of H3K27M-pDMG sEVs** (**A**) Percent of genes in sEVs from RR, RS, and RS-Clone 1 cells that are associated with the plasma membrane using the human proteome as a reference. (**B**) Top cellular components that represent the shared proteins among RR, RS, and RS-Clone 1 sEVs (**C**) - (**D**) Panther pathways that are represented by sEVs from RS, and RS-Clone 1 cells using a statistical overrepresentation analysis using a Fischer’s exact test. All pathway analysis was done using Panther. Cellular component analysis was done using G:profiler.

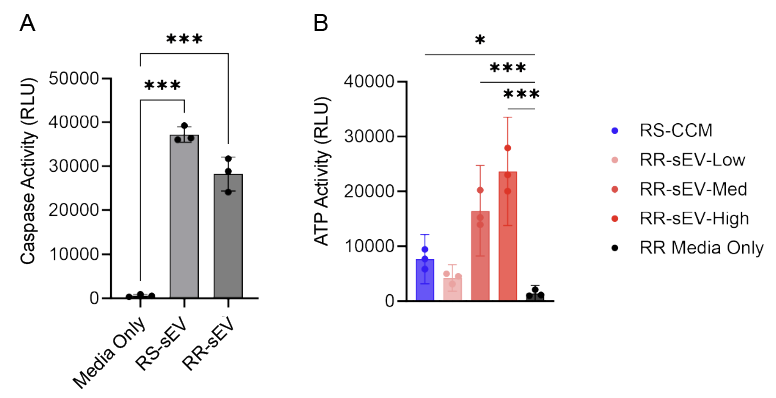

**Supplemental Figure 6: sEVs introduce confounding background signal in metabolic assays.** (**A**) Caspase-Glo 3/7 assay measuring luminescent caspase activity in media alone and after the addition of RS-sEVs or RR-sEVs. Both vesicle types induced high caspase signals (relative luminescence units, RLU) in the absence of cells, indicating potential assay interference. (**B**) CellTiter-Glo assay measuring ATP levels in media supplemented with increasing doses of RR-sEVs, RS-CCM, or RR media alone. RR-sEVs produced dose-dependent luminescent signal in the absence of cells, highlighting the limitations of this method for quantifying cell viability in the presence of sEVs. All experiments were performed in biological triplicate with at least two technical replicates per condition. Data are presented as means ± SEM. *p < 0.01, ***p < 0.0001 by one-way ANOVA with Tukey’s post-hoc test.

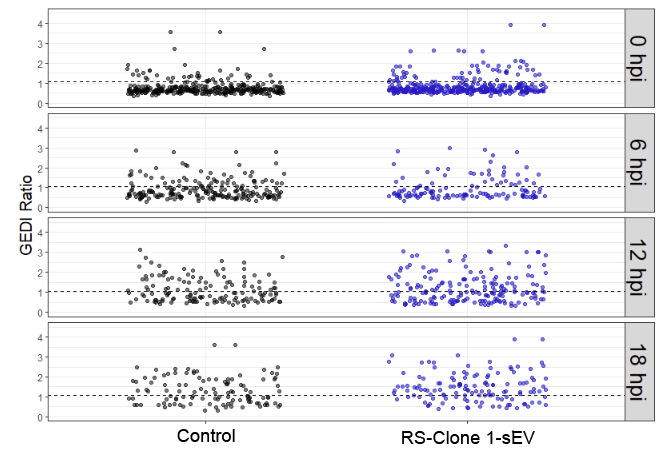

**Supplemental Figure 7: RS-Clone 1-sEVs improve survival of irradiated RS cells.** GEDI ratio values from RS cells stably expressing the GEDI construct, treated with RS-Clone 1-sEVs or control for 18 hours, followed by 8 Gy irradiation and live-cell imaging over 18 hours. Decreases in GEDI signal over time reflect death and detachment of cells from the plate. The experiment was performed with a seeding density of 2,500 cells per well, with at least two technical replicates per condition.
